# Redundant information across functionally coupled cortical networks supports rapid perceptual decisions in the ferret

**DOI:** 10.64898/2026.08.30.748127

**Authors:** Loren Koçillari, Edgar E. Galindo-Leon, Florian Pieper, Stefano Panzeri, Andreas K. Engel

## Abstract

Coordinated activity across cortical areas transforms sensory inputs into perceptual decisions, yet how task-relevant information is distributed across sites and linked to functional interactions and behavior remains unclear. Conventional functional connectivity measures reveal statistical dependencies between neural signals but cannot distinguish information encoded uniquely at individual sites, shared redundantly across sites, or available only from their joint activity. Here, we used Partial Information Decomposition (PID) to characterize stimulus information during fast and slow correct decisions. We analyzed local field potentials (LFPs) extracted from mesoscale electrocorticographic recordings from auditory, visual, and parietal cortices in ferrets performing a visual and audiovisual spatial-detection task. Time– and frequency-resolved analyses of local field potential power and phase showed that stimulus-side information was strongest in the theta and alpha bands and greater during fast than slow responses. PID applied to pairs of recording sites revealed that fast responses were associated with earlier and stronger unique information and a greater relative contribution of redundancy, whereas synergistic contributions were smaller. During fast responses, redundancy was selectively associated with stronger LFP power-envelope coupling. These findings indicate that faster perceptual decisions involve a frequency-specific reorganization of cortical information, characterized by early local encoding and enhanced redundant information across functionally interacting sites.

## Introduction

During everyday perception, coordinated activity across the mammalian cortex transforms sensory inputs into behavioral decisions. Previous work, including information-theoretic and decoding approaches, has shown that task-relevant information is distributed across multiple cortical areas (Delis et al., 2022; Hernández et al., 2010; International Brain Laboratory et al., 2025; Khilkevich et al., 2024; Lam et al., 2025; Lemke et al., 2024; Michalareas et al., 2016; Park et al., 2018; Siegel et al., 2015; Steinmetz et al., 2019; Wilming et al., 2020) and frequency bands, from theta to gamma (Bastos et al., 2015a; Belitski et al., 2010, 2008; Giraud and Poeppel, 2012; Gross et al., 2013; Kayser et al., 2009; Liebe et al., 2012). Recent studies have further shown that task-relevant variables can be encoded at the network level through pairwise and higher-order redundant and synergistic interactions among cortical areas (Combrisson et al., 2025, 2024; Gelens et al., 2024; Greco et al., 2024; Roberts et al., 2026). Together, these findings indicate that task-relevant information is organized not only locally but also distributed through the activity of many individual areas in different frequency bands and the interactions among cortical areas.

Functional connectivity provides a well-established framework for characterizing these interactions and relating them to behavior (Averbeck et al., 2006; Cohen and Maunsell, 2009; Engel et al., 2001; Engel and Gerloff, 2022; Fries, 2015; Gray et al., 1989; Panzeri et al., 2022; Pesaran et al., 2008; Siegel et al., 2012). Electrophysiological studies have typically distinguished two major forms of functional coupling: envelope coupling, which reflects slow co-modulations in signal power, and phase coupling, which quantifies the stability of inter-site phase relationships (Engel et al., 2013; Hipp et al., 2012; Lachaux et al., 1999; Nolte et al., 2004). Although these measures identify statistical dependencies between neural signals recorded at different locations, they do not determine how task-relevant information is distributed across recording sites. For example, measuring functional connectivity does not determine whether task-relevant information is carried independently by individual sites, redundantly shared across sites, or available only from their joint activity.

To characterize these information components, Partial Information Decomposition (PID) provides a framework for determining how task-relevant information is distributed across multiple recording sites. For a pair of sites, PID can be used to partition their joint information about a stimulus variable into unique, redundant, and synergistic components (Williams and Beer, 2010). Unique information is provided by one site but not the other; redundant information can be obtained from either site independently; and synergistic information refers to the additional information about the stimulus variable that becomes available only when the two sites are considered jointly. Redundancy and synergy have been associated with robustness and integration, respectively (Luppi et al., 2024b, 2024a), making PID well suited to characterizing distinct forms of neural information processing.

Previous studies have mainly characterized redundancy and synergy between neural activities recorded at different sites during rest, but have not measured synergy or redundancy of task-related information (Luppi et al., 2024b, 2024a). Other studies have examined redundancy and synergy of information about sensory stimulus variables during passive stimulus encoding, but have not assessed their behavioral relevance for task performance (Greco et al., 2024; Roberts et al., 2026). Another study examined redundancy and synergy of information gain during goal-directed learning, but did not relate these measures to behavioral performance (Combrisson et al., 2025). As a consequence, the relationship between redundancy and synergy and behavioral performance during active perceptual decision-making remains less well understood. Studies directly addressing this relationship have largely focused on local cortical circuits (Francis et al., 2022; Koçillari et al., 2023) or on pairs of areas (Lemke et al., 2024). Moreover, how synergy and redundancy are distributed across frequency bands during perceptual behavior remains poorly understood. It therefore remains unknown how behaviorally relevant redundancy and synergy are distributed across frequency bands and cortical sites spanning multiple areas at the mesoscale, and how these information components relate to frequency-specific functional connectivity.

Here, we address these questions using local field potentials (LFPs) obtained by low-pass filtering mesoscale electrocorticographic (ECoG) recordings from auditory, visual, and parietal cortices in ferrets performing a visual and audiovisual spatial-detection task (Galindo-Leon et al., 2025a; Hollensteiner et al., 2015). We first quantify frequency– and time-resolved information about a task-relevant stimulus feature—stimulus side—carried by LFP power and phase at individual recording sites, and determine how this information varies with response speed among correct trials. We then examine pairs of cortical sites by quantifying their joint stimulus information and applying PID to separate this information into unique, redundant, and synergistic components. Finally, we relate these information components to frequency-resolved functional connectivity, using amplitude-based envelope coupling for LFP power and phase-based coupling for LFP phase. This approach allows us to characterize how information associated with behavioral performance is organized across frequencies and cortical areas, and how it relates to frequency-resolved functional connectivity during perceptual decision-making.

## Results

### Stimulus-evoked LFP power and phase modulations

We analyzed cortical activity recorded with a chronically implanted 64-channel electrocorticographic (ECoG) array in ferrets performing a spatial detection task (see Methods) (Galindo-Leon et al., 2025a; Hollensteiner et al., 2015). The ECoG array covered auditory, visual and posterior parietal areas in the left hemisphere (Fig. 1A). Ferrets were well-trained on a two-alternative forced-choice spatial detection task in which they oriented their head toward a visual (V) or audiovisual (AV) stimulus presented on either the left or right side of an LCD monitor positioned in front of them. Hereafter, we refer to left and right stimuli as ipsilateral and contralateral, respectively, relative to the ECoG array. Stimulus side (contralateral vs. ipsilateral) therefore constituted the task-relevant stimulus feature (see Methods).

**Figure 1.**
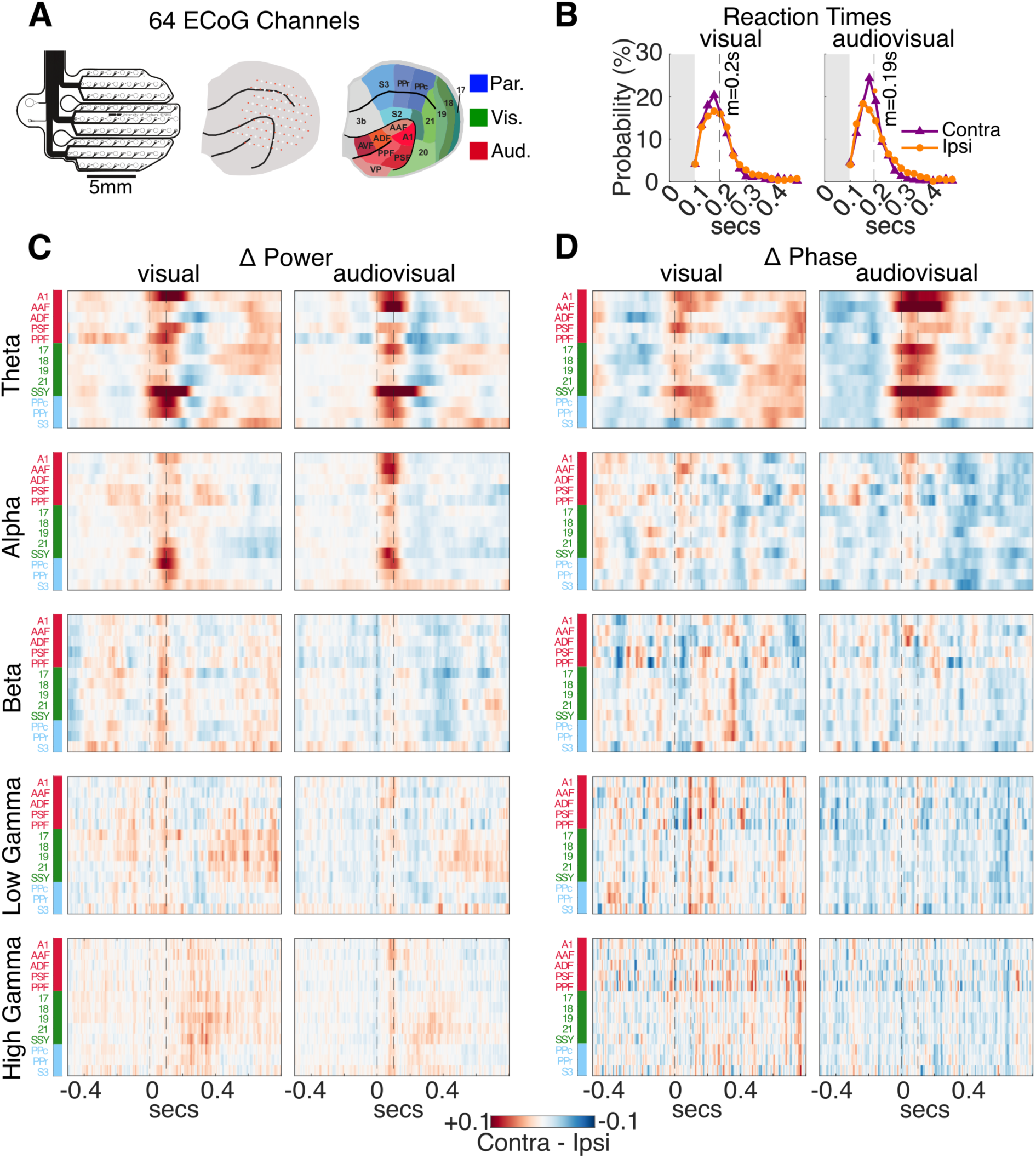
LFP power and phase time-courses during the lateralized detection task. **(A)** Schematic of the 64-channel ECoG array (left) implanted over the posterior cortex of the left hemisphere (middle) in awake, behaving ferrets, covering auditory (red), visual (green), and somatosensory/parietal (blue) regions (right), as defined by the functional parcellation of Bizley et al. (2007). **(B)** Reaction time (RT) distributions for ipsilateral (orange) and contralateral (purple) hit trials pooled across sessions and animals; vertical dashed lines indicate median RTs (0.20 s for visual and 0.19 s for audiovisual stimuli). **(C–D)** Time courses of contralateral–ipsilateral differences in LFP power **(C)** and phase **(D)** across cortical areas and frequency bands, computed across all hit trials. Vertical dashed lines denote stimulus onset and offset. Frequency bands: theta (4–8 Hz), alpha (8–16 Hz), beta (16–32 Hz), low gamma (32–64 Hz), and high gamma (64–128 Hz).

We first investigated whether behavioral reaction times (RTs), defined as the latency from stimulus onset to the first lick, differed by stimulus side. We restricted the analysis to hit trials because miss trials were substantially fewer in number (means±SD across subjects: 291±80 for visual and 509±216 for audiovisual hits; 57±17 for visual and 56±24 for audiovisual misses). RTs were analyzed separately for ipsilateral and contralateral trials and pooled across sessions and animals to form RT distributions. Median RTs were 200 ms for visual trials and 190 ms for audiovisual trials. RT distributions did not differ significantly between ipsilateral and contralateral stimuli for either visual or audiovisual conditions (two-sample Kolmogorov–Smirnov test: p = 0.5 for V trials; p = 0.28, for AV trials; Fig. 1B), indicating comparable reaction times across stimulus sides.

We then studied cortical activity evoked by the lateralized stimuli by analyzing LFP power and phase across time, frequency, and cortical regions. Channels were assigned to cortical regions according to the parcellation scheme of Bizley and coworkers (Bizley et al., 2007) (Fig. 1A). Median LFP power was first computed for each ECoG channel across trials separately for ipsilateral and contralateral conditions and then averaged across sessions and channels within each cortical area within each animal. Furthermore, we quantified inter-trial phase consistency (ITPC; see Methods) in ipsilateral and contralateral trials by computing circular means across trials, followed by averaging across sessions and channels within each cortical area for each animal. Finally, contralateral–ipsilateral differences in both mean power and ITPC were computed within each animal and subsequently averaged across animals (Fig. 1C–D).

Mean power and mean phase consistency were systematically larger in magnitude for contralateral than ipsilateral trials in theta– (4–8 Hz) and alpha-band (8–16 Hz) during the 200-ms post-stimulus interval (Fig. 1C-D). Higher-frequency bands showed only weaker and noisier contralateral-ipsilateral differences. LFP power in the visual condition peaked in the suprasylvian visual cortex (SSY) and posterior parietal caudal area (PPc) in the theta and alpha bands, and additionally in primary auditory cortex (A1) and posterior parietal rostral area (PPr) in the theta-band. In the audiovisual condition, LFP power peaked in auditory area AAF and SSY in the theta-band, and additionally in A1 and PPc in the alpha-band. Similarly, phase consistency was higher for contralateral than ipsilateral trials predominantly in the theta-band during audiovisual stimuli across A1, AAF, visual areas 17, 19, and SSY, whereas in the visual condition there was only a mild difference between contralateral and ipsilateral in A1, PSF, and SSY. Side differences in alpha-band phase consistency were much weaker than those in the theta-band and did not show clear patterns (Fig. 1D).

Together, these results demonstrate stimulus-related changes in power and in phase consistency across cortical regions, primarily in the theta– and alpha-bands. While these stimulus-related changes reveal average contralateral–ipsilateral differences across trials, they do not directly quantify how much information about stimulus side is carried by neural activity at individual recording sites. We therefore used information theory to quantify stimulus information at the single-channel level.

### Low-frequency LFP power and phase encode stimulus-side information

To quantify single-channel stimulus information, we computed Shannon’s mutual information (MI) (Cover and Thomas, 2006; Quian Quiroga and Panzeri, 2009; Shannon, 1948) about stimulus side (contralateral vs. ipsilateral) carried by either LFP power or phase. For each subject, MI was estimated using all available trials pooled across sessions and computed separately for each ECoG channel, frequency band, and time point.

The stimulus information, averaged across channels and subjects, was largest in the theta and alpha bands for both LFP power and phase (Fig. 2A). The beta, low-gamma, and high-gamma bands showed lower amounts of information. Thus, low-frequency LFP power and phase are particularly informative (Fig. 1C-D). We thus focused our subsequent analyses on the theta and alpha bands.

**Figure 2.**
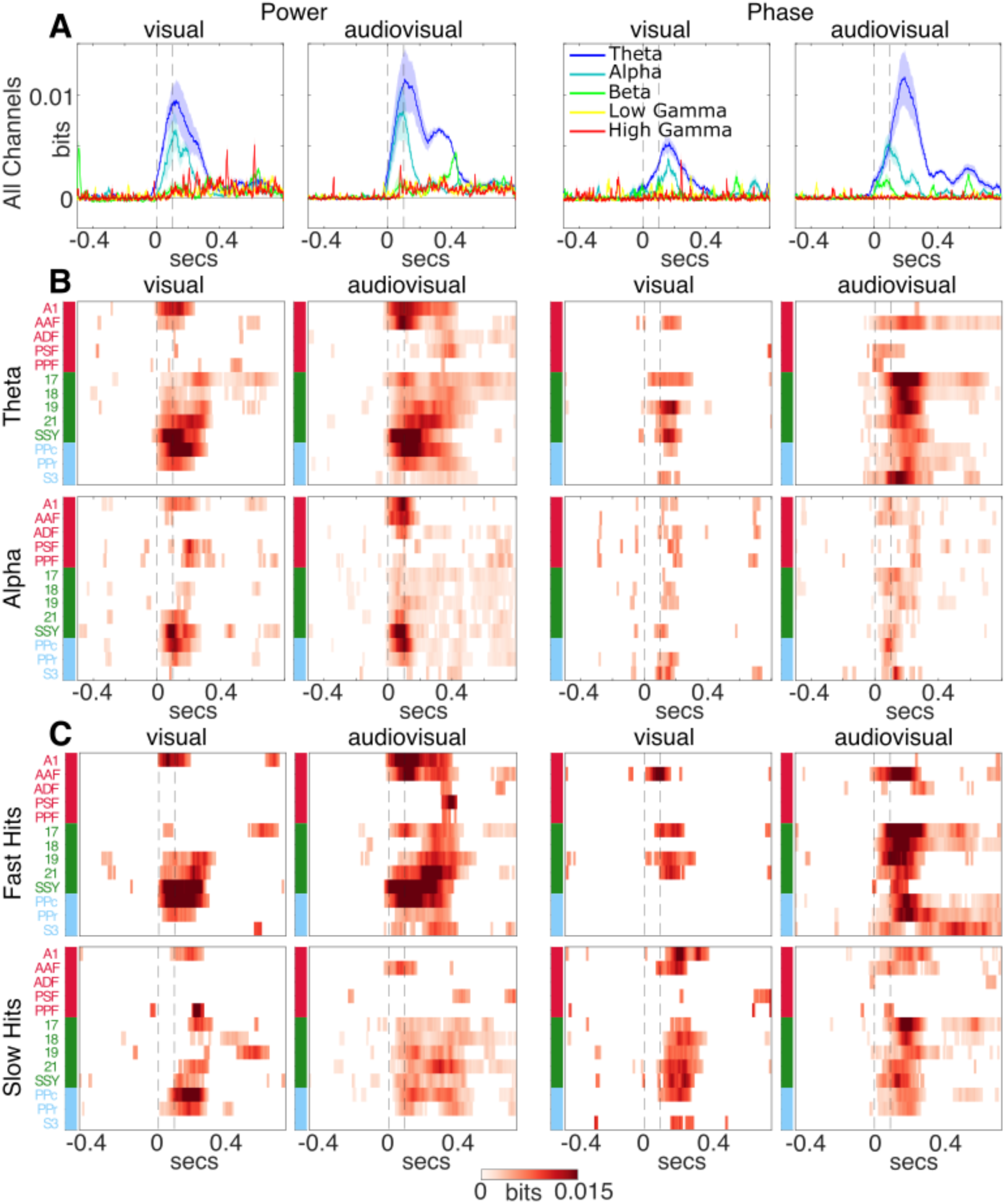
Stimulus-side information across time, cortical areas, and frequency bands. **(A)** Single-channel stimulus information time courses (mean ± SEM) averaged across all 64 ECoG channels for each frequency band. Information values significantly above the pre-stimulus baseline were identified by z-scoring single-channel time courses relative to the pre-stimulus baseline and applying Benjamini–Hochberg FDR correction across time (q ≤ 0.05); non-significant time points were set to zero. Pre-stimulus baseline information was subtracted from the entire information time course. **(B)** Cortical distribution of stimulus information in the theta– (top) and alpha– (bottom) bands, obtained by averaging time courses across channels within each brain area and then across animals. **(C)** Theta-band stimulus information computed separately for fast and slow hit trials, with trial categorization based on median reaction times for each stimulus modality (Fig. 1B). Vertical dashed lines denote stimulus onset and offset.

Theta-band information was sustained over a 400-ms post-stimulus interval, whereas alpha-band information persisted for a shorter ∼300-ms interval. In the theta-band, information peaked at approximately 120 ms for visual trials and 100 ms for audiovisual trials for LFP power, and at 140 ms for visual trials and 180 ms for audiovisual trials for LFP phase. In the alpha-band, information peaked at 110 ms for visual trials and 90 ms for audiovisual trials for LFP power, and at 170 ms for visual trials and 80 ms for audiovisual trials for LFP phase. Thus, across modalities and signal features, alpha-band information peaked earlier than theta-band information. However, theta-band information was consistently larger overall. We therefore focus mainly on theta-band results in the following, and report most of the alpha-band analyses in Supplementary Materials.

We next examined the cortical distribution of stimulus information in the theta– and alpha-bands (Fig. 2B). In the theta-band, stimulus information carried by LFP power during visual trials was strongest in SSY, PPc, and A1, with earlier peak latencies in SSY and A1 compared with PPc. During audiovisual trials, stimulus information additionally increased in the auditory area AAF. LFP phase conveyed less stimulus information than LFP power during visual trials showing a peak of information in visual areas 17 and 19. During audiovisual trials, stimulus information carried by LFP phase was larger in visual areas 17, 18, and 19 and in the somatosensory area S3. In contrast, alpha-band stimulus information carried by LFP power was generally weaker than in the theta-band but showed similar spatial patterns, with peaks of information observed in visual areas 21 and SSY, and in PPc for the visual condition; for the audiovisual condition, information additionally increased in A1 and AAF (Fig. 2B, bottom). For LFP phase, alpha-band information maps were generally weak and did not resemble the theta-band distribution, with no prominent peaks in the single-channel information time courses (Fig. 2B, bottom).

Finally, we tested whether stimulus information differed between rapid and slow correct responses (Fig. 2C). Within each stimulus modality, hits were categorized as fast or slow using the median reaction time (Fig. 1B). Across both visual and audiovisual modalities, stimulus information was higher in fast than for slow hits in both the theta-band (Fig. 2C) and the alpha-band (Fig. S1). This indicates that stimulus-related information is behaviorally relevant, with stronger stimulus encoding accompanying faster perceptual decision-making.

In summary, low-frequency LFP activity in the theta– and alpha-bands carried the largest amount of stimulus-side information. Stimulus information peaked shortly after stimulus offset and was enhanced during fast correct responses.

### Low-frequency information carried jointly by channel pairs and its decomposition into redundant, synergistic and unique information

Having characterized single-channel stimulus information and its relation to rapid and slow correct responses, we next asked how stimulus information in the low frequency bands that carry most information (theta and alpha band) is represented jointly by pairs of ECoG channels and how it differs between fast and slow hits. For each pair of channels, we computed the joint mutual information between stimulus side (contralateral versus ipsilateral) and the pair’s LFP power or phase time-courses (Fig. 3). We focused on channel pairs exhibiting statistically significant joint information, assessed with a non-parametric permutation test in which stimulus labels were randomized across trials and empirical peak joint-information values were compared against information peaks obtained from the resulting surrogate datasets (Francis et al., 2022). Multiple comparisons across pairs of channels were controlled using an FDR correction (q = 0.05) (see Methods).

**Figure 3.**
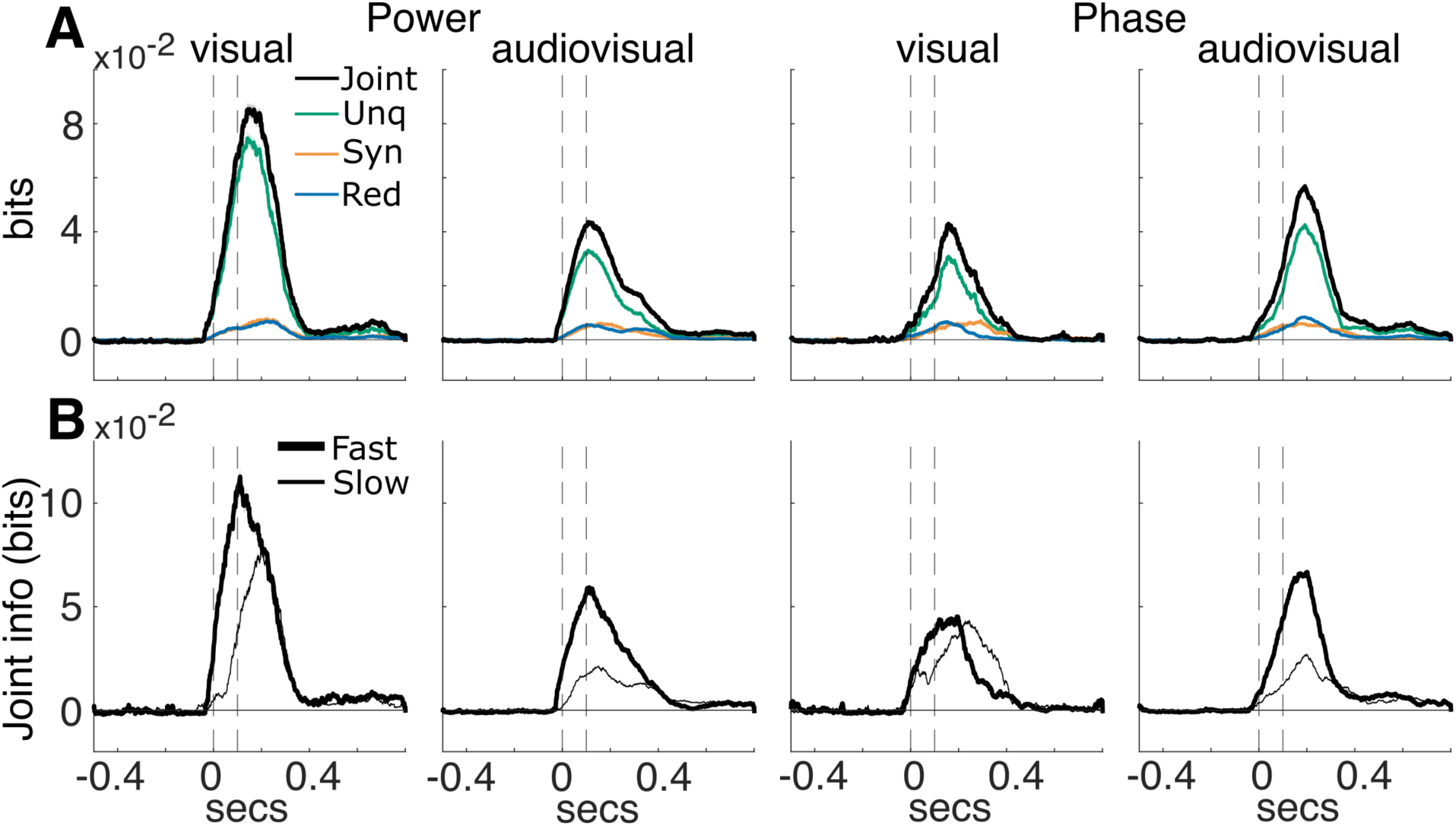
Time courses of theta-band joint information and its decomposition. **(A)** Time courses of theta-band joint information (black) and its unique (green), synergistic (orange), and redundant (blue) components, shown as mean ± SEM. Information was computed across all hit trials and averaged across all channel pairs carrying significant joint information. Significant joint information was identified using a non-parametric permutation test, as described in (Francis et al., 2022), followed by Benjamini–Hochberg FDR correction across channel pairs within each subject (q ≤ 0.05, see Methods). For each significant pair, information values significantly above the pre-stimulus baseline were identified by z-scoring relative to the pre-stimulus baseline and applying Benjamini–Hochberg FDR correction across time (q ≤ 0.05); non-significant time points were set to zero, and pre-stimulus baseline information was subtracted. **(B)** Joint information time courses computed separately for fast (bold lines) and slow (thin lines) hit trials. Shaded areas denote mean ± SEM across significant channel pairs pooled across subjects. Vertical dashed lines indicate stimulus onset and offset.

We first computed the joint information across all hit trials. Consistent with the single channel results, joint stimulus information carried by LFP power was higher for visual than for audiovisual stimuli, whereas LFP phase showed more information in audiovisual than visual stimuli (Fig. 3A). Joint information persisted for up to ∼400 ms in theta band and up to ∼200 ms in the alpha band after stimulus onset (Fig. S2). Moreover, we observed higher joint information in the theta-band than in the alpha-band (Fig. S3).

While joint information quantifies how much stimulus information is carried collectively by two channels, it does not reveal how that information is distributed across channels. To address this question, we applied the PID (Williams and Beer, 2010) to decompose joint stimulus information into unique, redundant, and synergistic information components. Unique information denotes stimulus information available exclusively in one channel but not the other, redundant information corresponds to information available in either channel, and synergistic information reflects information that emerges only when two channels are considered jointly. PID time courses were computed for all pairs of channels carrying significant joint information. The stimulus side was treated as the target variable whereas the two channels’ LFP power (or phase) as source variables. We focus the discussion and main figures on the theta band, which exhibited the highest single-channel information values. Unless otherwise stated, throughout the rest of the paper we therefore report joint information and its decomposition in the theta band. Results for the alpha band, including joint information and its components, are presented in the Supplementary Materials because they were more variable and less interpretable owing to their overall lower information values (Fig. S3).

Across conditions, unique information dominated the joint stimulus information (Fig. 3A), with the highest values observed for LFP power during visual trials. Although redundancy and synergy were consistently smaller in magnitude than unique information, both increased in the post-stimulus compared to the pre-stimulus interval, suggesting significant stimulus-related modulation. Statistical significance was assessed by comparing post-stimulus information values with those in the pre-stimulus interval using the FDR-corrected procedure as described in the Methods.

Together, these results show that joint stimulus information is primarily driven by unique contributions from individual channels. At the same time, redundant and synergistic components also increase after stimulus onset, suggesting that all PID information components contribute to stimulus-side processing.

### The behavioral relevance of joint information

To assess the possible behavioral relevance of the joint information and its components carried by LPF activity for rapid decision-making, we concentrated on differences in information during the first 150 ms after stimulus onset. We reasoned that if information values were higher and had shorter latency during this window for fast compared to slow hits, then these higher information values were relevant to support fast correct decisions. We therefore identified, within this behavioral-relevance window, information features that were selectively enhanced in fast-hit trials compared with slow-hit trials and considered them candidate information components most likely to support faster decisions. The 150-ms window was chosen based on the timing of behavioral responses. Because the median reaction time used to split hit responses into fast and slow responses was approximately 200 ms after stimulus onset for both stimuli (Fig. 1B), information components contributing to rapid perceptual decisions would need to become available early enough to influence the decision process before motor execution. We therefore focused the analysis to the first 150 ms after stimulus onset, assuming that ∼50 ms are needed before the behavioral response for motor-command generation and initiation of the overt movement.

We first computed joint information separately for fast and slow hit trials and consistently found larger values for fast than for slow hit trials in both the theta– (Fig. 3B) and alpha bands (Fig. S2B), including within the 150-ms behaviorally relevant window. In the theta-band, for LFP power, joint information reached a maximum of 0.11 bits in fast hits and 0.075 bits in slow hits during visual stimuli (Fig. 3B). During audiovisual trials, maximum information reached 0.056 bits in fast and 0.021 bits in slow hits. For LFP phase, joint information during visual stimulus reached a maximum of 0.045 bits in fast hits and 0.042 bits in slow hits, whereas during audiovisual stimuli it reached 0.067 bits in fast and 0.025 bits in slow hits.

In summary, joint stimulus information was consistently enhanced during fast compared with slow correct responses, including within the first 150 ms after stimulus onset, which suggests that early stimulus encoding of pairs of ECoG channels may contribute to faster correct decisions. We next decomposed this joint information to determine whether its behavioral relevance was primarily associated with unique, redundant, or synergistic information.

### The behavioral relevance of unique information and its cortical distribution

We first focused on the unique information. Across conditions, unique information was the largest component of the joint stimulus information (Fig. 3A), with the highest values observed for LFP power during visual trials. As reasoned above, we evaluated the behavioral relevant of unique information considering its timing and value during the behaviorally relevant window comprising the first 150 ms after stimulus onset.

Across channels belonging to pairs with significant joint information, unique information within the 150-ms behaviorally relevant window was, on average, larger in magnitude and peaked earlier for fast than for slow hits in both the theta (Fig. 4A) and alpha bands (Fig. S2B). This suggests that unique information is behaviorally relevant for rapid correct responses, with larger and earlier information associated with faster decisions.

**Figure 4.**
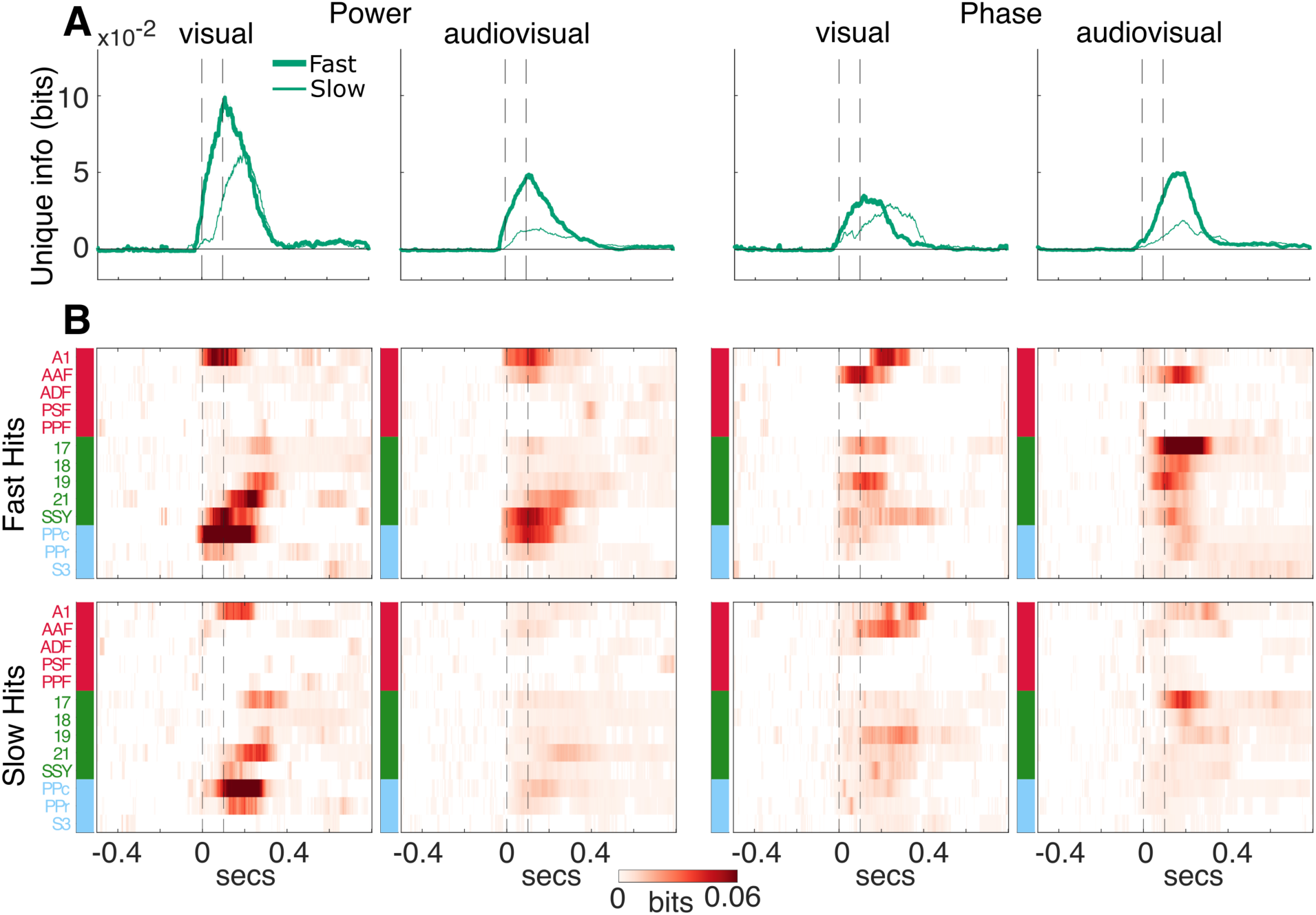
Cortical distribution of theta-band unique information during fast and slow hit trials. **(A)** Average unique-information time courses (mean ± SEM) across all channels belonging to channel pairs with significant joint information, computed separately for fast and slow hits. **(B)** For each brain area, time courses of unique information from channels belonging to significant channel pairs (carrying significant joint information) were pooled across animals and averaged. Information values significantly above the pre-stimulus baseline were identified by z-scoring each time course relative to its pre-stimulus baseline and applying Benjamini–Hochberg FDR correction across time (q ≤ 0.05); non-significant samples were set to zero, and pre-stimulus baseline information was subtracted. Vertical dashed lines indicate stimulus onset and offset.

We next characterized cortical distribution of unique information and asked whether its spatiotemporal dynamics differed between fast and slow perceptual judgments. For LFP power, unique information in fast correct trials was strongest in PPc, A1, and in visual areas SSY and 21 both in the theta-band (Fig. 4B) and alpha-band (Fig. S4) and for both visual and audiovisual stimulation. For visual trials, unique-information peaks were higher and occurred earlier in fast than in slow hit trials. For audiovisual trials, unique information likewise markedly decreased from fast to slow hits, indicating a strong relation of single-source stimulus representations to response speed.

Unique information also showed a distinct temporal organization across areas in fast compared to slow correct hits. During visual stimulation, unique information carried by LFP power exhibited a sequential pattern of information during the 150-ms behaviorally relevant window, with earlier onset in PPc and later onset in A1 and SSY followed by area 21. During audiovisual stimulation, peak latencies were more closely aligned across A1, SSY, and PPc, consistent with more temporally coordinated engagement of sensory and association regions when auditory and visual cues are combined.

For LFP phase, theta-band unique information for visual stimuli was highest in visual areas 17, 19, and SSY, as well as in the auditory area A1 and AAF, and was consistently larger in fast than in slow hits. For audiovisual stimuli, unique information was highest in visual areas 17, 18, 19, and SSY, as well as in the auditory area AAF, and was again consistently larger in fast than in slow hits. In contrast, alpha-band phase unique information was substantially weaker than theta– band phase information across both visual and audiovisual conditions (Fig. S3).

In summary, unique information carried by both LFP power and phase was larger and peaked earlier during the 150-ms behaviorally relevant window in fast compared to slow hits in both visual and audiovisual conditions and for both phase and power, showing a rapid onset in power in higher-order areas both during visual and audiovisual stimulation.

### The behavioral relevance of redundant, synergistic, and co-information

After characterizing differences in unique information between fast and slow hit trials, we next investigated redundant and synergistic information within channels pairs carrying significant joint information. We assessed the potential behavioral relevance of these components by comparing their timing and magnitude between fast and slow hits within the 150-ms behaviorally relevant window.

Within this window, both redundancy and synergy had consistently lower latency and reached higher peak values in fast than in slow hits, with the exception of the phase in the visual task, for which values were comparable between fast and slow hits (Fig.5A). These results suggest that, for LFP power, both redundant and synergistic information have a high behavioral relevance for taking fast correction decisions. For LFP phase, evidence for the behavioral relevance of redundancy and synergy was also present, but was less consistent than for power.

**Figure 5.**
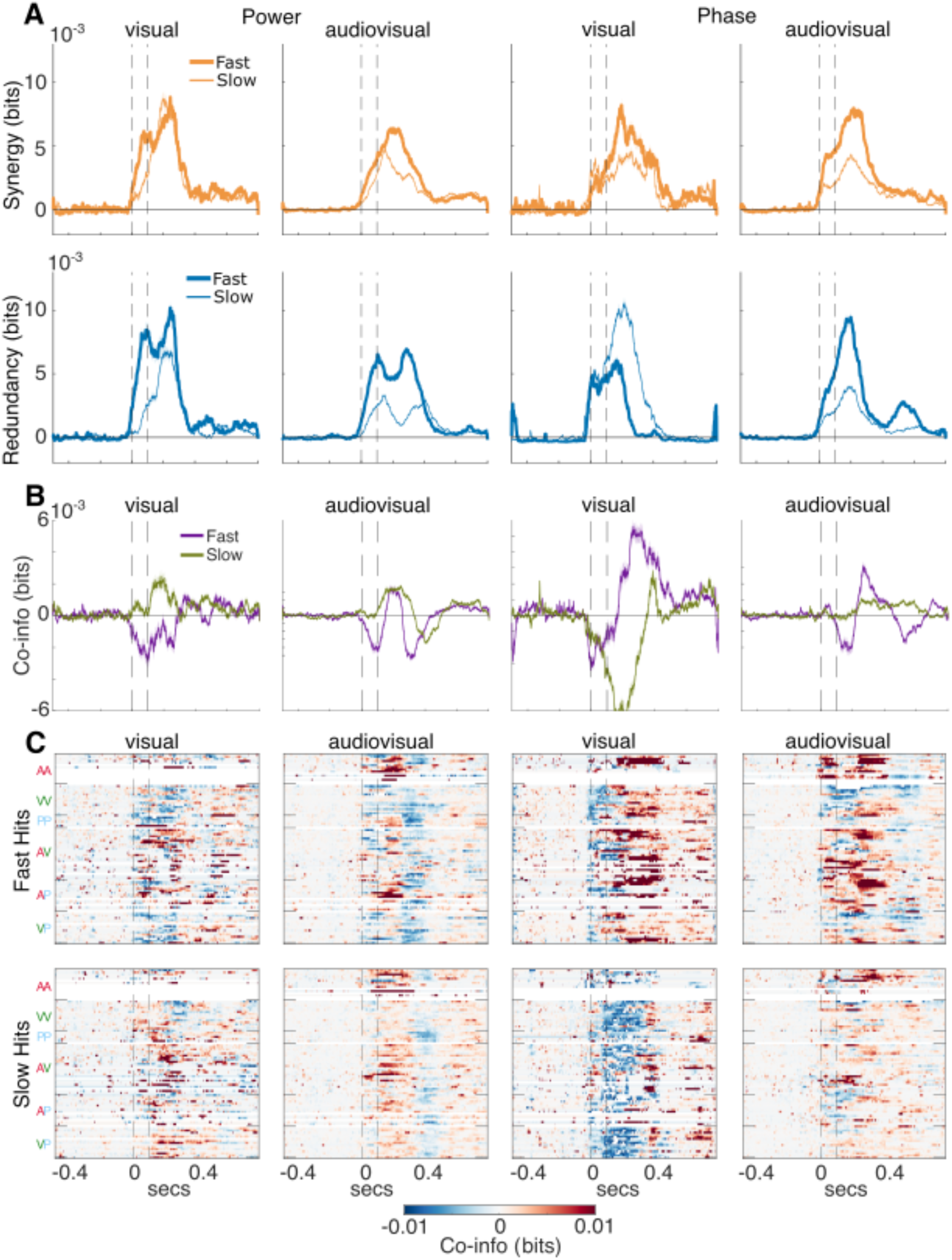
Theta-band redundant, synergistic, and co-information dynamics during fast and slow hit trials. **(A)** Average synergistic and redundant information and **(B)** co-information (synergy minus redundancy) time courses (mean ± SEM) across all channel pairs carrying significant joint information, computed separately for fast and slow hits. **(C)** Heatmaps showing co-information averaged across significant channel pairs and subjects for each pair of brain areas, grouped by functional system (A, auditory; V, visual; P, parietal). Synergistic and redundant values significantly above the pre-stimulus baseline were identified by z-scoring relative to the pre-stimulus baseline and applying Benjamini–Hochberg FDR correction (q ≤ 0.05); non-significant samples were set to zero. Vertical dashed lines denote stimulus onset and offset. Vertical dashed lines indicate stimulus onset and offset.

Another question was whether the balance between synergy and redundancy was behaviorally relevant. To quantify this balance, we next quantified co-information, defined here as synergy minus redundancy (Fig. 5B). Positive co-information values indicate synergy-dominated information, whereas negative values indicate redundancy-dominated information.

We studied how co-information differed between fast and slow hits. For LFP power, fast hits consistently showed negative co-information for both visual and audiovisual stimuli, whereas this pattern was not observed in slow hits, suggesting that a greater contribution of redundancy relative to synergy is associated with faster correct decisions (Fig. 5B). For LFP phase, during the 150-ms behaviorally-relevant window, co-information was negative only for fast hits in the audiovisual condition, whereas it was negative for both fast and slow hits for the visual condition. Thus, a predominance of redundancy over synergy may also be behaviorally relevant for phase information, although the evidence is less consistent than for power. Together, these results indicate that rapid correct responses are associated with stronger redundancy relative to synergy in theta-band LFP power.

To better elucidate which pairs of areas mainly contributed to the co-information profiles, we mapped the anatomical distribution of co-information (Fig. 5C), redundancy and synergy (Fig. S5) in the theta-band. Redundancy and synergy generally showed highly distributed topographies, involving channel pairs within and across multiple functional systems for both neurophysiological parameters. Redundancy of information in LFP power was prominent between parietal channels, between visual channels and in visual-parietal channel pairs during visual stimulation in fast trials (Fig. S5A). It showed an even more widespread involvement of most areas in the late phase of the trials in the audiovisual stimulation condition. Redundancy of information in the LFP phase was particularly widespread and prevailed across all areas and systems for slow hits in the visual condition (Fig. S5A, bottom panels). Synergy of information in LFP power was enhanced between parietal channels, as well as in audio-visual and visuo-parietal channel pairs in the visual condition (Fig. S5B). In the audiovisual condition, synergy of information in power occurred mainly between auditory channels, and in audio-visual channel pairs. Synergy of information in LFP phase strongly dominated during the late phase of fast trials, involving most areas with a widespread topography (Fig. S5B, bottom panels).

Overall, these results suggest that unique, redundant, and synergistic information are all behaviorally relevant for fast decision-making. Unique information within each cortical area appears to make the largest contribution to fast hit responses. The presence of both redundancy and synergy across channel pairs, with redundancy generally exceeding synergy, may provide an additional complementary contribution to rapid decisions, particularly for information carried by LFP power.

### Relationship of co-information to functional connectivity in fast and slow hits

Finally, we asked whether synergistic and redundant information relate to large-scale functional connectivity (FC), quantified using amplitude-based (envelope) coupling for LFP power and phase-based coupling for LFP phase(Engel et al., 2013), and whether these relationships differ between fast and slow hit trials.

We restricted the analysis to pairs of channels carrying statistically significant joint information and quantified FC over time using a sliding-window approach (100-ms windows advanced in 50-ms steps; Fig. 6A; see Methods). Surrogate-based statistics were used to classify channel pairs as significantly connected or non-significantly connected within each window (see Methods). Repeating the analysis with longer windows (500-ms) yielded qualitatively similar results (not shown).

**Figure 6.**
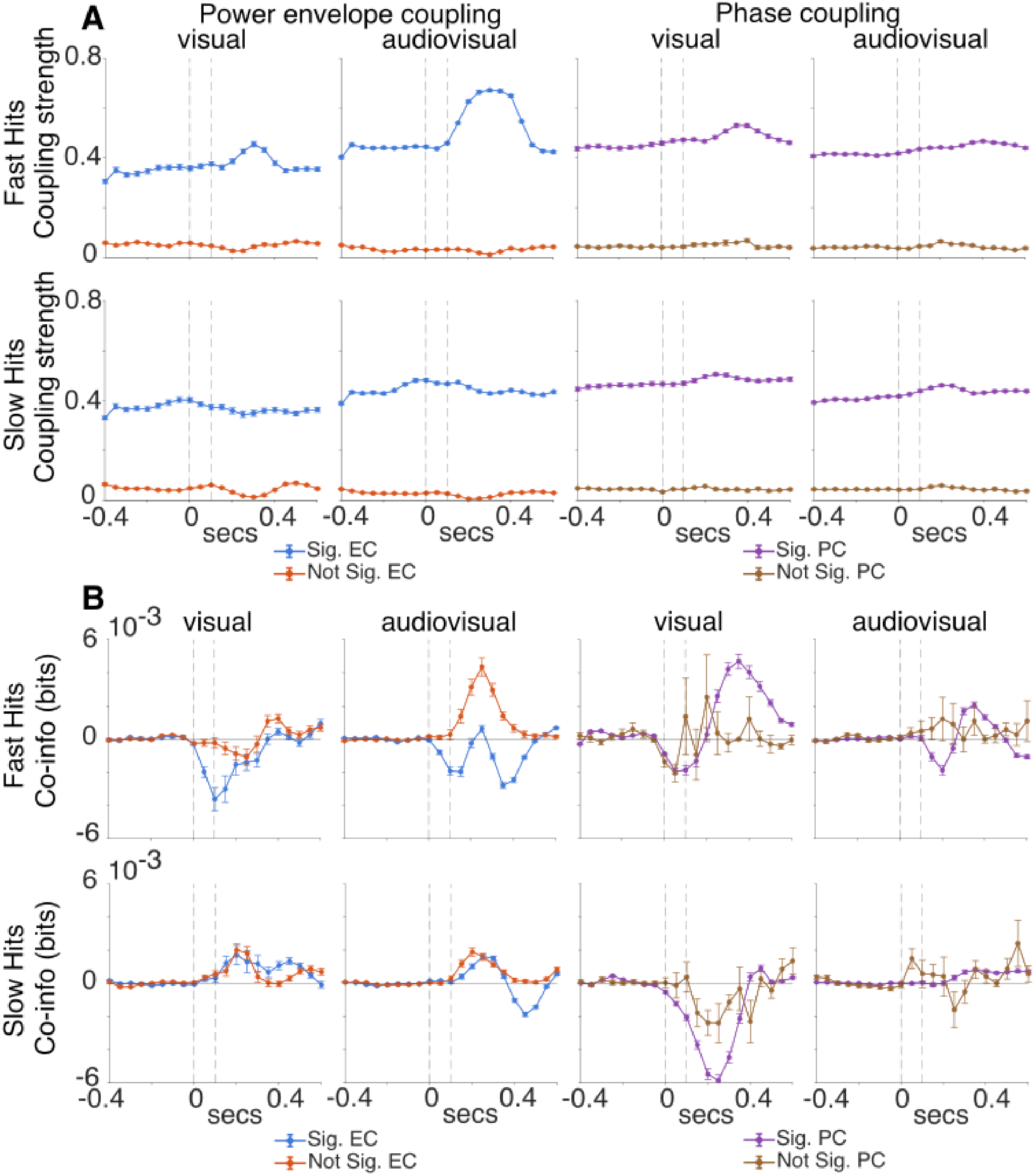
Theta-band co-information in functionally coupled channel pairs during fast and slow hits. **(A)** Time-courses of theta-band functional connectivity for channel pairs carrying significant joint information (mean ± SEM), computed using the envelope coupling (EC) and phase coupling modes (PC). We labelled each FC link as significant if they passed a statistical test (see Methods) and we labelled them as significant. **(B)** Time courses of theta-band co-information for channel pairs carrying significant joint information, computed separately for functionally connected (Sig FC) and non-connected pairs during fast and slow hits. Co-information was estimated using a sliding window of 100 ms advanced in 50 ms steps. Functional connectivity was assessed within each window using envelope coupling for LFP power and phase coupling for LFP phase, with significance determined by a circular-shift permutation test (p < 0.05). Statistical differences between conditions were evaluated using an unpaired t-test (**p < 0.001). Vertical dashed lines indicate stimulus onset and offset.

Amplitude envelope coupling in connected pairs exhibited a clear behavioral modulation: during fast trials it displayed a post-stimulus increase (∼200-400 ms) compared to the pre-stimulus interval, whereas slow hits did not show a comparable modulation (Fig. 6A). Phase coupling in connected pairs increased only weakly compared to pre-stimulus in fast hits and remained largely unchanged in slow hits. Across stimulus conditions, amplitude envelope coupling tended to increase more during audiovisual than visual condition, whereas phase coupling exhibited a stronger increase compared to the pre-stimulus interval during visual than audiovisual stimulation. The fraction of significantly connected pairs differed across coupling metrics (Fig. S7): for amplitude envelope coupling, connected and unconnected pairs were mode balanced (visual: ∼50/50; audiovisual: ∼70/30), whereas phase coupling classified most pairs as connected (∼95%) in both conditions. Together, these results suggest that amplitude-based coupling is more behaviorally modulated than phase coupling, which instead remains broadly similar across fast and slow hits.

We then examined how FC relates to theta-band co-information (Fig. 6B). We averaged co-information within each sliding window and compared its values in the two groups of significantly connected versus non-significantly connected pairs of channels. For amplitude envelope coupling, functionally connected channel pairs showed systematically negative co-information in fast hits, with a pronounced redundancy-dominant information peaking around ∼100 ms post-stimulus for visual and around ∼100 and ∼400 ms for audiovisual stimulus condition. This result was driven by higher redundancy levels in connected pairs (Fig. S8A), while synergy was comparable between connected and unconnected pairs for the visual stimulation, whereas it decreased in connected pairs in the audiovisual stimulation. In contrast, slow hits exhibited a trend for positive co-information across stimulus modalities, indicating a shift toward synergy (Fig. 6B, bottom panels). However, for slow hits connected pairs of areas showed both, higher synergy (Fig. S8A) and higher redundancy (Fig. S8B) than not connected pairs. In the alpha-band, predominance of redundancy was also observed for fast hits and, to a lesser degree, for slow hits (Fig. S9).

For phase coupling, in fast hits, co-information was negative in functionally connected pairs during the stimulus interval (0-100 ms) in the visual condition and during the 100-200 ms interval in the audiovisual stimulation. In both stimulation conditions, co-information changed to positive values in the later part of the trials (Fig. 6B, top panels). As shown in Fig. S8, redundancy and synergy were both increased for connected pairs in fast hits, with a slightly later increase of synergy compared to redundancy (Fig. S8A, B, top panels). Non-significantly connected pairs showed increases in both synergy (Fig. S8A) and redundancy (Fig. S8B) during and after the stimulus intervals. In the slow hits, for visual stimuli, co-information was negative in both connected and unconnected pairs of channels, but it was more negative in connected pairs indicating a higher level of redundancy (Fig. 6B, bottom panels; Fig. S8A, B, bottom panels). In the audiovisual stimulation, co-information was around zero, suggesting more balanced values of synergy and redundancy. Overall, our results indicate that phase coupling was associated with increases of both synergy and redundancy, whereby functionally connected pairs showed larger fluctuations in co-information than unconnected pairs (Fig. 6B).

To link these findings to the topography of functional connections defined by envelope and phase coupling, we examined how FC and PID components were distributed across pairs of cortical areas by constructing area-level matrices during the early stimulus interval (0–100 ms; Fig. S10) and the post-stimulus interval (300-400 ms; Fig. S11). PID components exhibited distinct spatial patterns between fast and slow hits which differed between the stimulus interval (Fig. S10) and the post-stimulus interval (Fig S11). During the stimulus interval of the visual condition, synergy in LFP power was higher in auditory-parietal and auditory-visual pairs of areas for fast hits compared to slow hits (Fig. S10). Redundancy in LFP power showed a similar effect, with additional increase in redundancy in visual-parietal pairs. Differences in synergy and redundancy for LFP phase between fast and slow hits were less pronounced during the stimulus interval (Fig. S10). Functional connectivity in the stimulus interval showed only mild differences between fast and slow hits, indicating that behavioral differences in PID components cannot be directly explained by changes in coupling strength alone.

During the post-stimulus interval, PID components difference as well as FC differences between fast and slow hits showed a qualitatively distinct picture (Fig. S11). In this interval, marked differences in amplitude envelope coupling could be observed for the audiovisual condition between fast and slow hits for most pairs of areas. Interestingly, the higher envelope coupling for fast hits was paralleled by higher redundancy across most systems (Fig. S11A). Differences in phase coupling were less pronounced between fast and slow hits. However, higher synergy for LFP phase could be observed for fast hits between visual and parietal, as well as auditory and parietal areas. (Fig. S11A). Interestingly, pairs of areas with the highest synergy for LFP phase were generally not identical to those with the strongest phase coupling. In contrast, redundancy for LFP phase was higher for slow hits in the cluster of visual areas in the visual stimulation condition (Fig. S11B).

Together, these results suggest that the synergy-redundancy balance differs between functionally connected and unconnected cortical areas, with the clearest distinction arising during fast perceptual decisions. Specifically, functionally connected areas showed an early relative increase in redundancy over synergy during fast hit trials, particularly among pairs coupled through the amplitude envelope. This suggests that envelope coupling may promote redundant coding of LFP power across cortical networks, thereby supporting rapid perceptual decisions. By contrast, unconnected areas showed relatively greater synergy or a more balanced synergy-redundancy profile. During slow hit trials, both connected and unconnected areas showed either synergy-dominated information or a more balanced ratio between synergy and redundancy, indicating that redundancy-dominated coding was not simply a marker of correct performance but was preferentially associated with faster responses, except for the visual condition, where phase coupling showed dominant redundancy in both connected and unconnected pairs.

## Discussion

Here, we decomposed task-related sensory information at the mesoscale in ferrets to determine how its components relate to behavioral performance and whether redundancy and synergy are linked to large-scale functional coupling. To address these questions, we combined frequency-resolved ECoG recordings with partial information decomposition (PID) to quantify unique, redundant, and synergistic information across time, frequency bands, and cortical areas in ferrets performing a visual and audiovisual spatial detection task. Using the reaction time distribution, we classified correct trials into fast and slow hits. Stimulus-related information encoded in LFP power and phase was higher in fast hits, was concentrated primarily in the theta-band, and was represented in both early sensory and higher-order areas. Our PID analysis revealed that rapid correct performance was associated with a combination of early unique information and the presence of both synergy and redundancy, with a relative predominance of redundant over synergistic coding during the pre-response interval, particularly for LFP power. Together, these patterns indicate that rapid perceptual processing benefits from the emergence of early unique information, alongside both redundant and synergistic information, with the balance shifted toward redundancy. These changes in PID components showed only a moderately strong relationship with functional connectivity quantified by amplitude envelope and phase coupling.

### Low-frequency stimulus information

A key finding of our study is that single-channel stimulus information was carried predominantly in the theta-band and, to a lesser extent, in the alpha-band, in both LFP power and phase across stimulus conditions. Moreover, stimulus-related information was higher in fast than in slow correct responses, suggesting that stronger sensory encoding supports faster decisions. This result is consistent with previous work showing that theta-band activity encodes stimulus information (Belitski et al., 2010, 2008; Montemurro et al., 2008; Panzeri et al., 2010), and is related to response speed (Louviot et al., 2025; Ye et al., 2023).

The cortical distribution and temporal profile of stimulus information differed between LFP power and phase. For LFP power, stimulus information emerged early in the trial and was prominent in higher-order areas, including posterior parietal cortex (PPc) and the suprasylvian visual cortex (SSY), as well as in the early sensory area A1. On the other hand, LFP phase showed higher and earlier information in early visual areas 17 and 19 than in PPc and PPr, although its information peak occurred later than that of LFP power. Our findings contrast with the classical hierarchical feedforward framework, in which sensory information is expected to propagate from early sensory areas (e.g., V1) to higher-order regions. Instead, the early engagement of higher-order regions in LFP power is more consistent with a contribution of top-down influences in addition to bottom-up sensory processing. The delayed phase-based information in early visual areas may likewise reflect modulation of bottom-up processes by earlier top-down processes observed in power. This interpretation is compatible with the demands of the present task, which implied mainly stimulus localization rather than fine feature discrimination or multisensory integration. Furthermore, the animals were highly trained to the detection task, which may favor rapid recruitment of higher-order circuits involved in evidence accumulation and response guidance.

Overall, these findings suggest that low-frequency dynamics, particularly in the theta-band, support rapid performance based on the sensory encoding in early sensory and higher-order cortical areas. One plausible interpretation is that greater theta-band information observed in fast hits reflects a task state in which attention or temporal expectation optimizes low-frequency dynamics to support rapid sensory processing, thereby enabling earlier downstream readout and shorter reaction times. This interpretation is consistent with evidence that theta-band activity contributes to attentional gain control and in the temporal organization of cortical excitability, which can shape how effectively sensory evidence is sampled and propagated through cortical hierarchies (Fiebelkorn and Kastner, 2019; Fries, 2015; Helfrich et al., 2018; Schroeder and Lakatos, 2009). However, this interpretation remains speculative, as the available data did not include independent behavioral or physiological measures of the animals’ attentional state.

### Behavioral roles of the information components

PID analyses revealed that theta-band joint information was dominated by unique information across stimulus conditions and signal features, including both LFP power and phase. Although synergistic and redundant components were smaller in magnitude, both components contributed to stimulus encoding. Importantly, each component was also modulated by decision speed, suggesting that successful performance depends not only on how much information is represented, but also on how that information is distributed across cortical areas.

Unique information emerged earlier and was larger in fast than in slow hits, and its cortical distribution closely mirrored that of single-channel information. For LFP power, unique information was strongest in SSY, PPc, and A1, whereas for LFP phase it was strongest in early sensory processing stages, including A1, AAF, and visual areas 17 and 19. These findings further support the interpretation that rapid correct decisions rely on both early bottom-up processing, reflected more strongly in LFP phase, and the engagement of higher-order circuits, reflected more strongly in LFP power. In this context, local unique information may provide an early and spatially distributed substrate for rapid perceptual readout, potentially shaped by attentional or expectation-related state influences.

Redundancy and synergy were likewise behaviorally modulated, but in distinct ways. Across modalities, fast hits were characterized by a greater relative contribution of redundant than synergistic information in LFP power, whereas this contribution was reduced in slow hits, suggesting that redundancy in power is associated with faster responses. Because redundant information emerged earlier and was stronger than synergy in most conditions, it is likely to be more directly relevant for rapid behavioral responses in the present task. This speed-dependent modulation challenges the view that redundancy is merely an information-limiting byproduct. Whereas classical efficient-coding theories emphasized redundancy reduction as a key principle of efficient sensory representation (Attneave, 1954; Barlow, 1961; Laughlin, 1981; Simoncelli and Olshausen, 2001), our findings are consistent with recent studies highlighting the functional benefits of redundancy, such as improved robustness and reliability of downstream sensory readout (Francis et al., 2022; Koçillari et al., 2023; Panzeri et al., 2022; Valente et al., 2021). From this perspective, redundancy may facilitate rapid decisions by ensuring that task-relevant information is shared across multiple sites, enabling robust readout even when individual channels are noisy or only partially informative.

Synergy played a comparatively smaller role than redundancy in the present task, possibly because the task placed limited demands on integration across areas. This interpretation leads to the prediction that paradigms involving stronger ambiguity, greater working-memory demands, or multi-stage evidence accumulation may amplify synergistic contributions and reveal a more direct behavioral role for synergy.

### Relation to functional coupling modes

A largely unexplored topic is the relation of redundancy and synergy with classical measures of functional connectivity (FC) at mesoscales (coupling modes). While previous studies have linked redundancy and synergy to functional connectivity at the microscale (Koçillari et al., 2023) and macroscale (Luppi et al., 2022; Pope et al., 2025), it has been less explored how these information components relate to frequency-specific functional coupling during behavior. Here, we focused on two commonly studied coupling modes: amplitude-based coupling, quantified by envelope correlation, and phase-based coupling, quantified by the phase-locking value (Engel et al., 2013; Hipp et al., 2012; Lachaux et al., 1999). These low-frequency coupling modes have been linked to complementary functional roles, with amplitude envelope coupling often interpreted as reflecting slower co-fluctuations in population activity and large-scale functional network organization (Brookes et al., 2011; Engel et al., 2013; Hipp et al., 2012), and phase coupling as reflecting more precise temporal coordination and effective inter-areal communication (Bastos et al., 2015b; Fries, 2015; Womelsdorf et al., 2006).

Our results reveal a substantial dissociation between coupling patterns and information components. The topographical distribution of FC networks in the theta-band was broadly similar across fast and slow hits, whereas the corresponding PID networks differed substantially. Thus, behaviorally relevant changes in information sharing cannot be explained by differences in connectivity strength alone, but instead reflect changes in how information is distributed across cortical networks embedded within similarly coupled functional networks. This interpretation is consistent with previous work suggesting that redundant and synergistic information architectures are not captured by traditional pairwise FC measures (Luppi et al., 2022; Varley et al., 2023b).

A more specific relationship between PID and FC emerged when considering whether area pairs were functionally connected. During the stimulus interval on fast correct choices, PID analyses exhibited larger relative redundancy in functionally connected than unconnected pairs for both envelope– and phase-coupling, with the strongest effect observed for envelope coupling. This finding suggests that functionally connected areas can carry overlapping stimulus information, consistent with a network regime that emphasizes shared and reliable representations. During slow hits, in contrast, the amounts of redundancy and synergy were more balanced. These findings align to previous studies showing that redundancy is selectively enriched among functionally connected pairs and is behaviorally relevant (Francis et al., 2022; Koçillari et al., 2023).

Taken together, these results support the view that interareal coupling promotes reliable information sharing by reinforcing common representations across distributed networks. The increased redundancy among connected pairs of cortical areas during fast correct responses further links functional connectivity to behaviorally relevant information processing. Together, these findings identify a role of large-scale functional coupling, particularly in the theta-band, in supporting rapid perceptual decision-making through robust redundant information transmission.

### Methodological considerations

Our study extends previous work on redundancy and synergy at rest (Luppi et al., 2022; Pope et al., 2025) and during passive stimulus encoding (Greco et al., 2024; Roberts et al., 2026) by linking PID components to a behaviorally relevant stimulus feature. Among task-related PID studies (Delis et al., 2022; Park et al., 2018; Varley et al., 2023c), we adopted a complementary approach by treating the task-relevant stimulus feature as the PID target variable and neural signals as sources, rather than using neural activity itself as the target. For instance, Park and colleagues (Park et al., 2018) reported beneficial theta-band synergistic and redundant interactions between auditory and visual stimuli for human speech perception, but framed neural activity as the PID target. By focusing on the stimulus as the target, our approach allows a more direct interpretation of how redundant and synergistic information in neural activity reflects behaviorally relevant sensory information. Moreover, whereas previous work applied PID to learning signals (e.g., information gain (Combrisson et al., 2025)), our study applied PID to a perceptual task using a behaviorally relevant stimulus feature, thereby enabling a direct characterization of redundant and synergistic sensory information supporting perceptual decisions. Finally, we extended recent task-related findings linking stimulus-related redundancy between pairs of auditory neurons to correct decisions in a mouse tone-discrimination task (Koçillari et al., 2023) by showing that, at the mesoscale across cortical areas, redundancy is enhanced during fast, correct perceptual decisions.

### Future directions

One of the key questions in cognitive neuroscience is to elucidate the physiological role of coupling modes in information processing (Engel et al., 2013; Engel and Gerloff, 2022). Although a substantial body of work has linked functional connectivity to cognition and sensorimotor processing, this literature has only established correlative associations between coupling modes and behavioral variables (Engel et al., 2013; Fries, 2015; Reid et al., 2019; Siegel et al., 2012), leaving unresolved whether functional connectivity plays a causal role in shaping information processing and cognition (Engel and Gerloff, 2022). The same limitation applies to the present findings: while we observed associations of redundancy, synergy, functional connectivity, and behavior, information-theoretic measures alone cannot establish causality. An important next step will therefore be to combine this framework with causal perturbations, such as optogenetic manipulations, electrical stimulation, or pharmacological interventions, to test whether redundancy-dominated information in functionally connected pairs contributes causally to rapid correct decisions.

A second open question concerns the distinct and potentially complementary roles of envelope and phase coupling in information processing (Engel and Gerloff, 2022; Galindo-Leon et al., 2025b, 2019). Current evidence suggests that these coupling modes are not redundant, especially during sensory stimulation (Nolte et al., 2020; Siems and Siegel, 2020). Recent work further indicates that they can interact during the resting-state dynamics (Galindo-Leon et al., 2025b) and exhibit distinct spatial topographies and stimulus dependencies (Galindo-Leon et al., 2019). What remains unresolved is whether, during task performance, amplitude envelope and phase coupling carry overlapping or complementary information about task variables, and whether their joint contributions to stimulus or choice encoding is additive, redundant, or synergistic. More specifically, it will be important to determine whether the two coupling modes convey distinct task-relevant information along different cortical pathways, and whether combining them improves prediction of task variables beyond either mode alone. Furthermore, our study was confined to PID analysis of theta-band activity, and it will be important to investigate information carried by coupling modes in other frequency bands. Addressing these questions by jointly quantifying the information carried by both coupling modes, and by explicitly testing their unique, redundant, and synergistic contributions, will be an important direction for future work.

Finally, the present findings should be corroborated across a broader range of task designs, including paradigms with greater cognitive complexity. The predominance of redundancy observed here may reflect the demands of a rapid spatial detection task involving transient stimuli and relatively low cognitive load. By contrast, more complex behaviors, such as those requiring working memory, flexible rule switching, or hierarchical inference, may rely more strongly on synergistic information (Proca et al., 2024). Extending this framework to a broader range of tasks and behavioral contexts will be crucial for understanding how unique, redundant, and synergistic information adapt to changing cognitive demands.

## Materials and Methods

### Behavioral task

We analyzed ECoG recordings previously collected in published studies (Galindo-Leon et al., 2025a; Hollensteiner et al., 2015). The full experimental procedures are described in detail in (Galindo-Leon et al., 2025a); here we provide a concise overview. Ferrets were trained on a lateralized detection task, responding to brief (100 ms) visual (V) or audiovisual (AV) stimuli presented to the left or right of the virtual vertical midline of a screen positioned in front of them. During the task, animals were head-free but body-fixed inside a cylindrical tube positioned in front of the screen. Each trial started once the animal maintained a stable, straight head position for 500 ms while a stationary random-noise pattern was displayed to in the center of the screen to guide eye-sight and sustain attention. The contrast of this pattern decreased, signaling the animal to expect a stimulus appearance within the next second. The stimulus appeared at a random time on either the left or right side of the screen. It was either a unimodal visual circular grating, or an audiovisual stimulus consisting of the same visual grating paired with a spatially congruent white-noise burst.

Animals were required to remain still throughout the stimulus window. A 700-ms response interval followed, during which ferrets indicated stimulus side by turning their head toward the left or right. Correct responses were rewarded with water and allowed immediate initiation of the next trial. Trials were terminated if the animal responded before stimulus onset or within 100 ms thereafter, selected the incorrect side, or failed to respond. Aborted trials were followed by a 2-s timeout before the next trial could begin. A staircase procedure was applied to control the animal’s behavior, adjusting the difficulty through contrast/amplitude changes to achieve an average performance of around 75% correct responses (Galindo-Leon et al., 2025a; Hollensteiner et al., 2015). To further aid control of performance, catch-trials without any stimulus appearance took place at random occasions, where the animal was simply rewarded for keeping its head in the center-position throughout the whole trial.

Each ECoG contact was assigned to a cortical area (visual, auditory, parietal, or somatosensory) by topographically mapping its location onto the ferret cortical parcellation of (Bizley et al., 2007). Electrode positions were reconstructed from photographs acquired during surgery, and channels were manually labeled to cortical areas for each animal.

### Electrophysiological data preprocessing

The dataset consists of 57 recording sessions from four animals. To maximize statistical power, trials were pooled across sessions separately for each animal. Raw ECoG signals were acquired with a 64-channel AlphaLab SnR system at either 44.6 or 44 kHz and converted to local field potentials (LFP) by downsampling to 1395.1 Hz in two subjects and to 1375 Hz in the other two subjects. Signals were band-pass filtered with a high-pass cutoff at 0.1 Hz and a low-pass cutoff at 1/4^th^ of the respective LFP sampling frequency. Continuous recordings were segmented into 1.3–s trials aligned to stimulus onset. Each trial comprised a 500-ms pre-stimulus baseline and an 800-ms post-stimulus interval that included the 100-ms stimulus period—during which animals were instructed to remain still—and the subsequent 700-ms response period.

Filtered signals were re-referenced on a trial-by-trial basis by subtracting, at each timepoint, the mean across all channels. For each channel, pre-stimulus activity was pooled across trials to compute baseline mean and standard deviation, which were then used to z-score the time series on each trial.

To study the neural dynamics across frequency bands, preprocessed LFPs were band-pass-filtered into five frequency bands: theta (4–8 Hz), alpha (8–16 Hz), beta (16–32 Hz), low gamma (32–64 Hz), and high gamma (64–128 Hz). Filtering was performed with a zero-phase, second-order Butterworth filter implemented in MATLAB: band-specific coefficients were obtained with *butter*, and signals were filtered forward and backward using *filtfilt* to avoid phase distortions.

Because LFP data were analyzed at slightly different sampling frequencies across animals, either 1395.1 Hz or 1375 Hz, we resampled each animal’s time-resolved LFP power, phase, and information time courses onto a common 1-ms temporal grid (1 kHz) for plotting purposes only, enabling point-by-point averaging across animals.

### LFP power and phase

To extract time-resolved LFP power and phase, we computed the analytic signal of each band-pass-filtered signal for every channel and trial using the MATLAB function *hilbert*. For a real-valued filtered signals *x*(*t*), the analytic signal is defined as 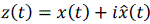, where 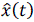 is the Hilbert transform of *x*(*t*) (Boashash, 1992):

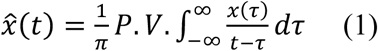

with P.V. denoting the Cauchy principal value. From the analytical signal *z*(*t*), we computed the instantaneous amplitude and phase. Instantaneous amplitude was defined as the modulus of *z*(*t*):

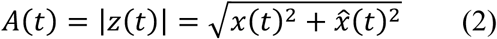

LFP power (P(t)) was defined as the square of the instantaneous amplitude:

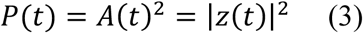

Instantaneous phase, here referred to as LFP phase, was computed as the argument of *z*(*t*) using the MATLAB function *angle*:

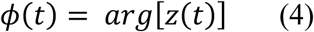

### Amplitude envelope and phase coupling

To quantify functional connectivity over time, we used a sliding-window approach (100-ms windows advanced in 50-ms steps). Within each window, LFP power or phase time series were concatenated across trials to form, for each channel, a single one-dimensional time series per condition (fast vs. slow hits). Functional connectivity was then computed within each window on these concatenated time series using an amplitude-based metric for LFP power (envelope coupling) and a phase-based metric for LFP phase (phase coupling). Amplitude-based functional connectivity (envelope coupling) was quantified as the Pearson correlation between LFP power time courses from pairs of channels within a given window:

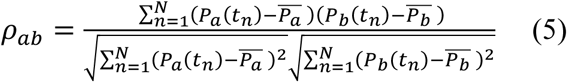

where *P_a_*(*t_n_*) and *P_b_*(*t_n_*) denote the LFP power at sample n for channels a and b, N is the number of samples in the window, and 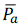 and 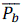 are the corresponding mean power values within that window.

Phase-based functional connectivity was quantified using the phase-locking value (PLV) (Lachaux et al., 1999), which measures the consistency of the instantaneous phase difference between two signals. For channels a and b at carrier frequency f, PLV was computed within each window as:

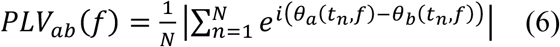

where *θ_a_*(*t_n_*, *f*) and *θ_b_*(*t_n_*, *f*) are the instantaneous phases at sample n, and the normalization by N constrains PLV to the range [0,1], with higher values indicating more consistent phase differences across time samples.

### Statistical significance of functional connectivity

To assess whether functional connectivity exceeded chance level, we used a circular-shift permutation test. For each pair of channels, one of the two concatenated time series was circularly shifted by a randomly selected lag, expressed as an integer number of samples, while the other time series was held fixed. This procedure preserves each signal’s autocorrelation structure while disrupting the temporal alignment. This procedure was repeated 100 times per pair to obtain a null distribution of coupling values (e.g., Pearson correlation for envelope coupling). A two-sided p-value (p-value<0.05) was then computed by comparing the absolute value of the observed coupling to the distribution of absolute surrogate values. Channel pairs whose coupling exceeded the surrogate distribution were labelled as significantly functionally connected.

### Computation of single-channel stimulus information

We quantified how much stimulus information the instantaneous LFP power and phase of each ECoG channel carried about stimulus side (contralateral vs. ipsilateral). Mutual information was computed separately for each stimulus modality (V, AV), frequency band, and time point using Shannon’s definition for two discrete random variables (Cover and Thomas, 2006; Quian Quiroga and Panzeri, 2009):

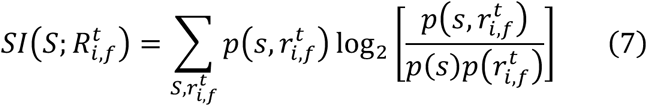

where S = (Contralateral, Ipsilateral) denotes stimulus category and 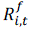 denotes the response of channel *i* at time point *t* and frequency band *f*. For each time point *t*, response distributions 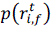 were estimated by discretizing responses across trials. LFP power values were binned into two equally populated bins. LFP phases values were first centered by subtracting the circular mean across trials and then discretized into two equally spaced bins corresponding to the two halves of the unit circle (equivalently, positive vs. negative imaginary component).

### Information above pre-stimulus baseline

To identify time points carrying stimulus information above the pre-stimulus, we normalized each information time course by z-scoring it relative to the pre-stimulus interval. Depending on the analysis, these time courses corresponded either to single-channel information, computed separately for each channel and each animal, or to PID time courses, computed for pairs of channels. This procedure assumes that pre-stimulus information values are approximately Gaussian distributed, allowing z-scores to be converted into p-values. Z-scores were converted to one-tailed p-values testing for increases above baseline, and multiple comparisons across time were controlled using the Benjamini–Hochberg false discovery rate (FDR) procedure (q ≤ 0.05). Time points surviving FDR correction retained their original values, whereas non-significant samples were set to zero. Finally, each time course was baseline-corrected by subtracting the mean pre-stimulus information, computed over the 500 ms interval preceding stimulus onset.

### Robustness analyses of information estimates: binning and temporal resolution

To test the robustness of information estimates to the discretization of response distributions, we repeated the analysis using larger number of bins (nbins = 4, 6, and 8). As expected, increasing the number of bins on the response variables increased absolute information values, but the temporal profiles of information and condition-dependent patterns remained highly similar (Fig. S13). We therefore used nbins = 2 in the main analyses to reduce estimation bias, particularly for joint mutual information and its decomposition into PID components, for which synergistic information is known to be particularly bias-sensitive (Lorenz et al., 2026).

We further evaluated how stimulus-information estimates depend on the temporal scale over which information is computed by using a sliding-window approach with window lengths ranging from ∼1 to 200 ms. For window lengths between ∼1 and 50 ms, information estimates were comparable to the single-time-point (∼1-ms) results shown in Fig. 2. For longer windows, we observed a frequency-dependent trade-off: gamma-band information increased while theta-band information decreased relative to estimates without temporal averaging (Fig. S12). For LFP phase, increasing the window size produced a consistent reduction in low-frequency stimulus information. Based on these observations, we retained single-time-point estimates to maximize temporal precision in subsequent analyses of both LFP power and phase.

### Computation of joint stimulus information and Partial Information Decomposition (PID)

We computed joint stimulus information as the mutual information between stimulus category S and the joint responses of two channels i and j, at time t in frequency band f:

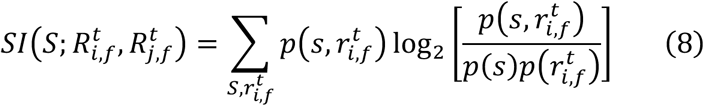

We then decomposed 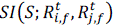 into non-negative information components—synergy, redundancy, and the two unique information encoded by each channel using the Partial information Decomposition (PID):

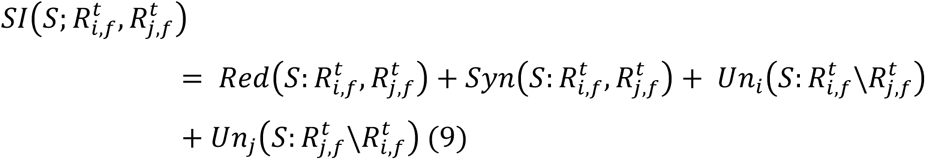

where 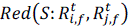 is redundant information present in both channels, 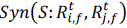 denotes the synergistic information that arises only when the two channels are considered jointly, and 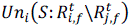 and 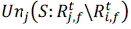 are the unique information contributions carried exclusively by channels i and j, respectively. Because PID components are linked by Shannon information identities, estimating one component (together with the relevant entropies) is sufficient to derive the remaining three components by linear combinations. To compute the PID components we used the approach outlined in (Bertschinger et al., 2014). Bertschinger et al. first defined the unique information in terms of a constrained convex optimization problem as follows:

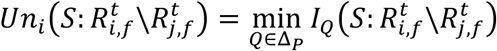

where the minimization occurs in the space Δ*_P_* of trivariate probability distributions 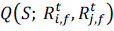 I with fixed marginals 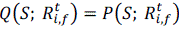 and 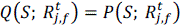. We numerically solved this problem using the MINT Toolbox (Lorenz et al., 2025), which implements the BROJA estimator (Bertschinger et al., 2014) after discretizing responses as in the single-channel analyses.

### Statistical significance of joint stimulus information

Statistical significance of joint stimulus information was assessed using a non-parametric permutation test (Francis et al., 2022). For each pair of channels, we compared the empirical peak mutual-information to a null distribution of peak values derived from surrogate datasets. Specifically, 100 surrogate datasets were generated by randomly shuffling stimulus labels across trials while keeping the LFP power or phase time series of both channels unchanged. This procedure preserves the marginal response distributions while destroying any stimulus–response dependency. A p-value was computed as the proportion of surrogate peak values that were greater than or equal to the empirical peak joint mutual information.

Pairs of channels with p < 0.05 were deemed significant, and only their information time courses were retained for subsequent analyses. We controlled for multiple comparisons across channel pairs using the Benjamini–Hochberg FDR procedure (q ≤ 0.05) applied to all 8,064 channel pairs (2,016 per subject, 4 subjects in total)). Significant pairs of channels were then selected for the subsequent PID analyses.

### PID and FC matrices

PID and FC values were grouped across pairs of ECoG channels assigned to each pair of cortical areas, and area-level matrices were constructed (one entry per area pair). Each matrix entry represents an average across channel pairs and then across animals.

### Software used

We developed custom software to analyze the data. For the computation of stimulus information and PID we used the MINT toolbox (Lorenz et al., 2025).

## Data and software availability

The dataset and software are available upon request.

## Acknowledgements

This work was supported by the European Research Council, project cICMs, ERC-2022-AdG-101097402 (awarded to A.K.E.). Views and opinions expressed in this paper are those of the authors only and do not necessarily reflect those of the European Union or the European Research Council. Neither the European Union nor the granting authority can be held responsible for them.

## Author contributions

L.K., A.K.E. and S.P. conceived and designed the study. L.K. performed the research, analyzed the data, and discussed the results with E.E.G.-L., A.K.E. and S.P. L.K., E.E.G.-L. and F.P. contributed to data-preprocessing. L.K. prepared the figures and wrote the first draft of the manuscript. A.K.E and S.P. supervised the study. A.K.E acquired funding. All authors reviewed and edited the manuscript, and approved the final version.

## Supplementary Figures

**Figure S1.**
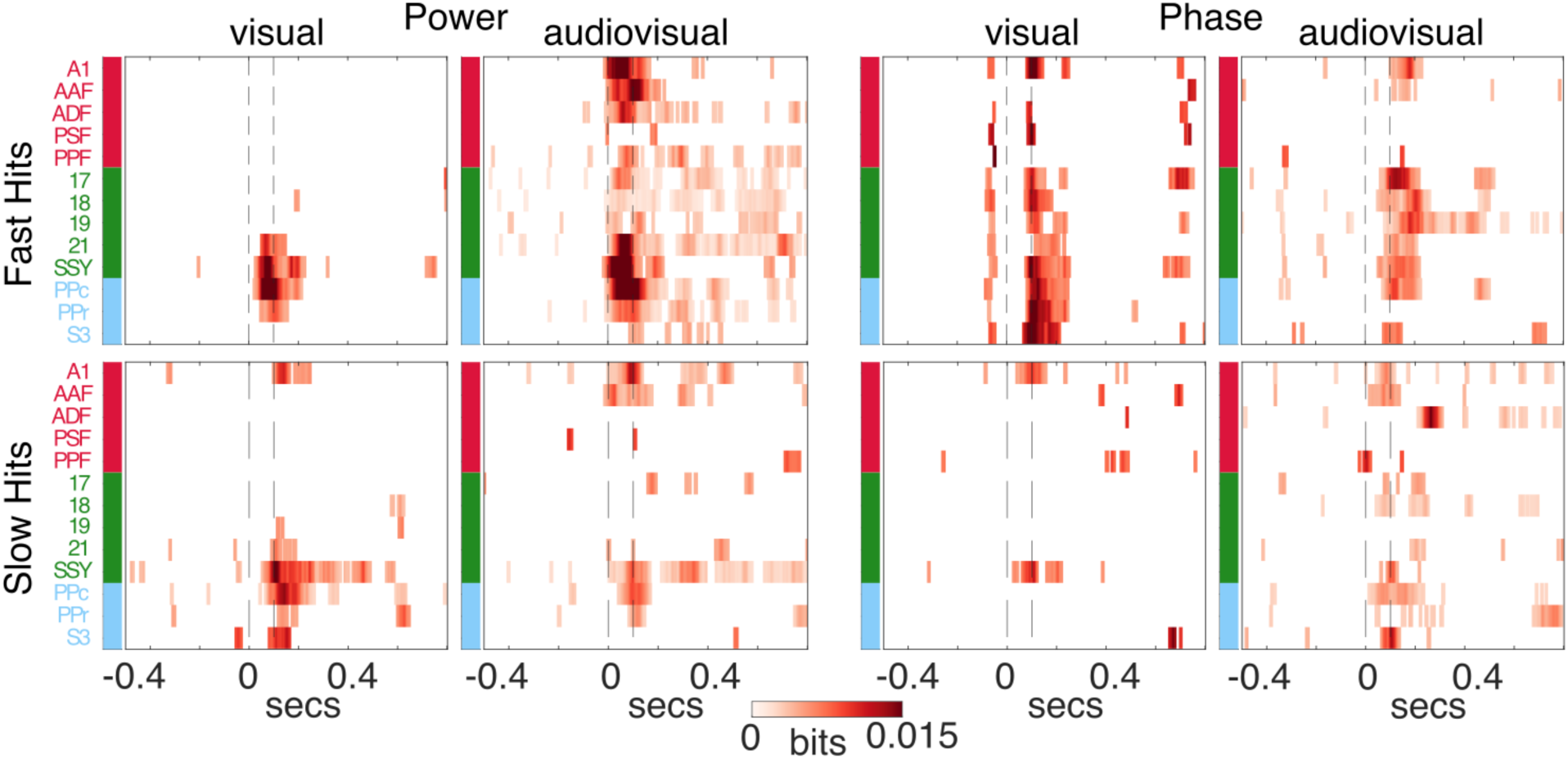
Alpha-band stimulus information during fast and slow hit trials. Alpha-band stimulus information time courses are shown separately for fast and slow hits. Significant time points were identified by z-scoring single-channel time courses relative to the pre-stimulus baseline and applying Benjamini–Hochberg FDR correction across time (q ≤ 0.05); non-significant samples were set to zero, and pre-stimulus baseline information was subtracted. Trial categorization into fast and slow hits was based on median reaction times for each stimulus modality (Fig. 1B). Vertical dashed lines denote stimulus onset and offset.

**Figure S2.**
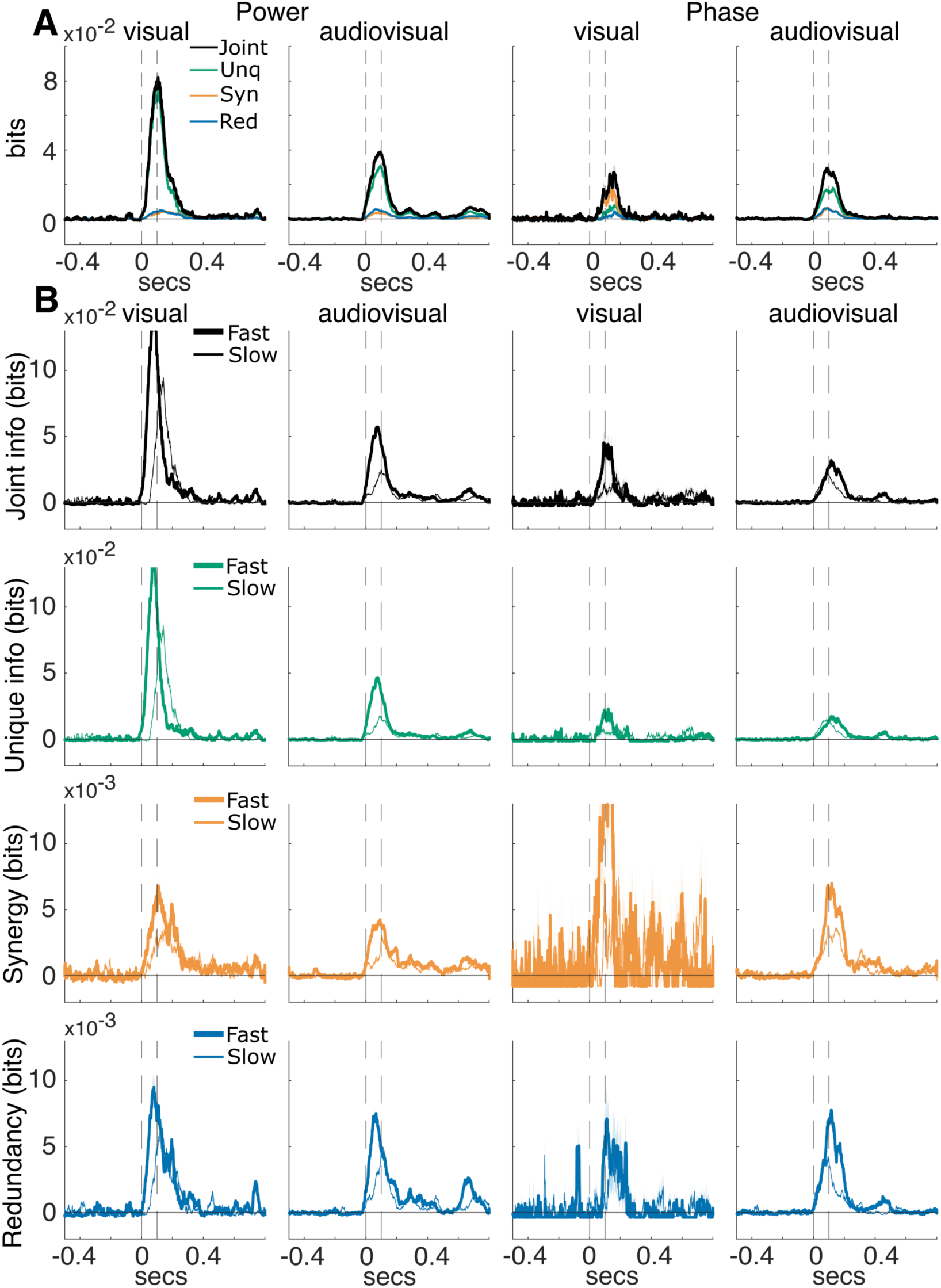
Time-courses of alpha-band joint, synergistic, redundant and unique information. **(A)** Time courses of alpha-band joint (black), unique (green), synergistic (orange), and redundant (blue) information (mean ± SEM), averaged across all channel pairs carrying significant joint information (identified by permutation testing with Benjamini–Hochberg FDR correction across channel pairs, q ≤ 0.05). For each significant pair, significant time points were identified by z-scoring relative to the pre-stimulus baseline and applying Benjamini–Hochberg FDR correction across time (q ≤ 0.05); non-significant samples were set to zero, and pre-stimulus baseline information was subtracted. **(B)** Corresponding information time courses computed separately for fast (bold lines) and slow (thin lines) hit trials. Shaded areas denote mean ± SEM across significant channel pairs pooled across subjects. Vertical dashed lines indicate stimulus onset and offset.

**Figure S3.**
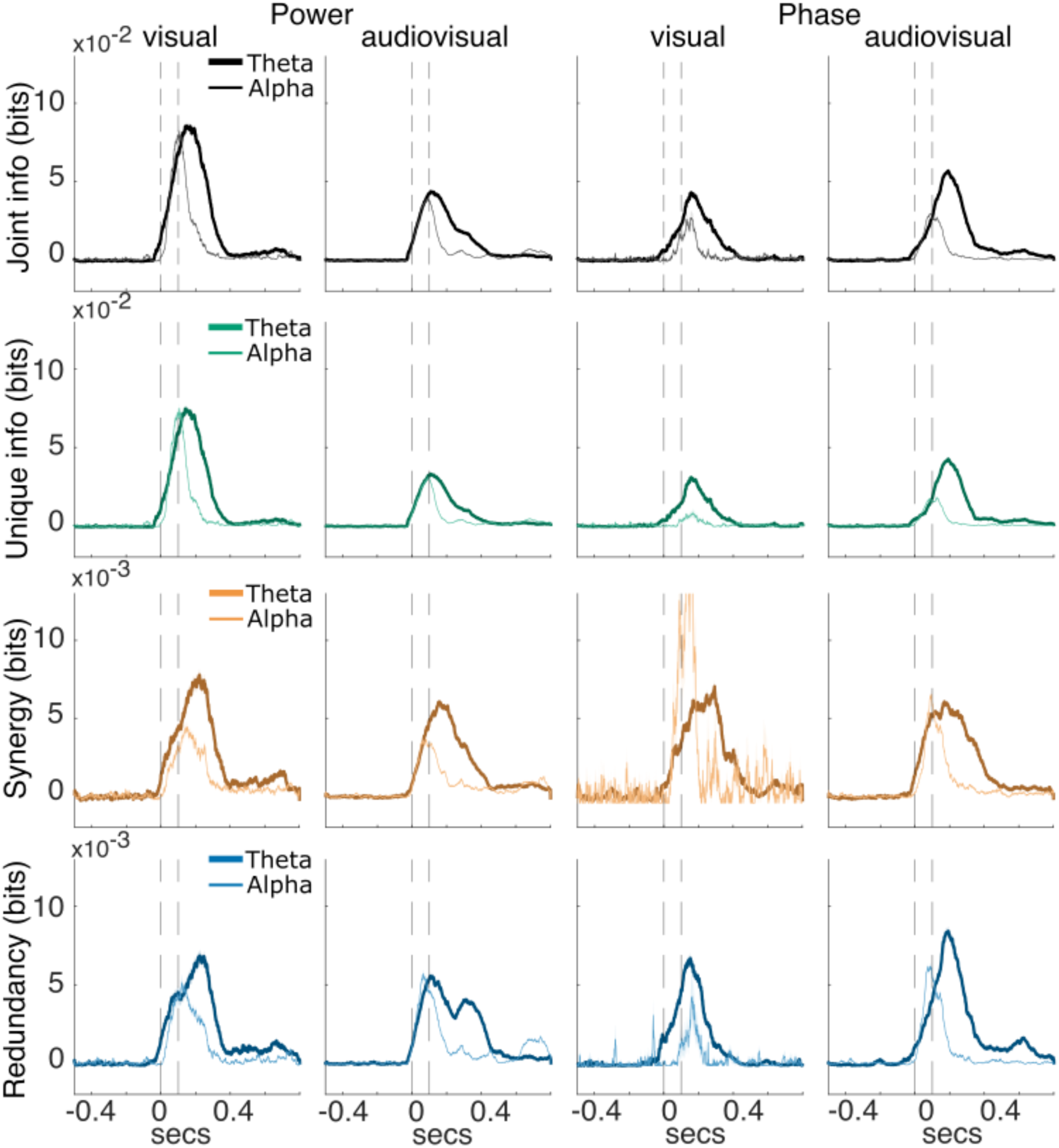
Time-courses of synergistic, redundant and unique information for theta and alpha bands for hit trials. Time courses of joint information (black), unique information (green), synergy (orange), and redundancy (blue), for theta (bold lines) and alpha (thin lines) bands computed for theta band (bold lines) and alpha band (thin lines) (mean ± SEM). Significant pairs of channels were identified based on significant joint information in theta and alpha bands, respectively (see Methods). For each information time-course we subtracted its average pre-stimulus information. Vertical dashed lines indicate stimulus onset and offset, respectively. Bold lines and shaded areas indicate the mean ± SEM across all significant channel pairs exhibiting significant joint information, pooled across subjects. Vertical dashed lines mark stimulus onset and offset.

**Figure S4.**
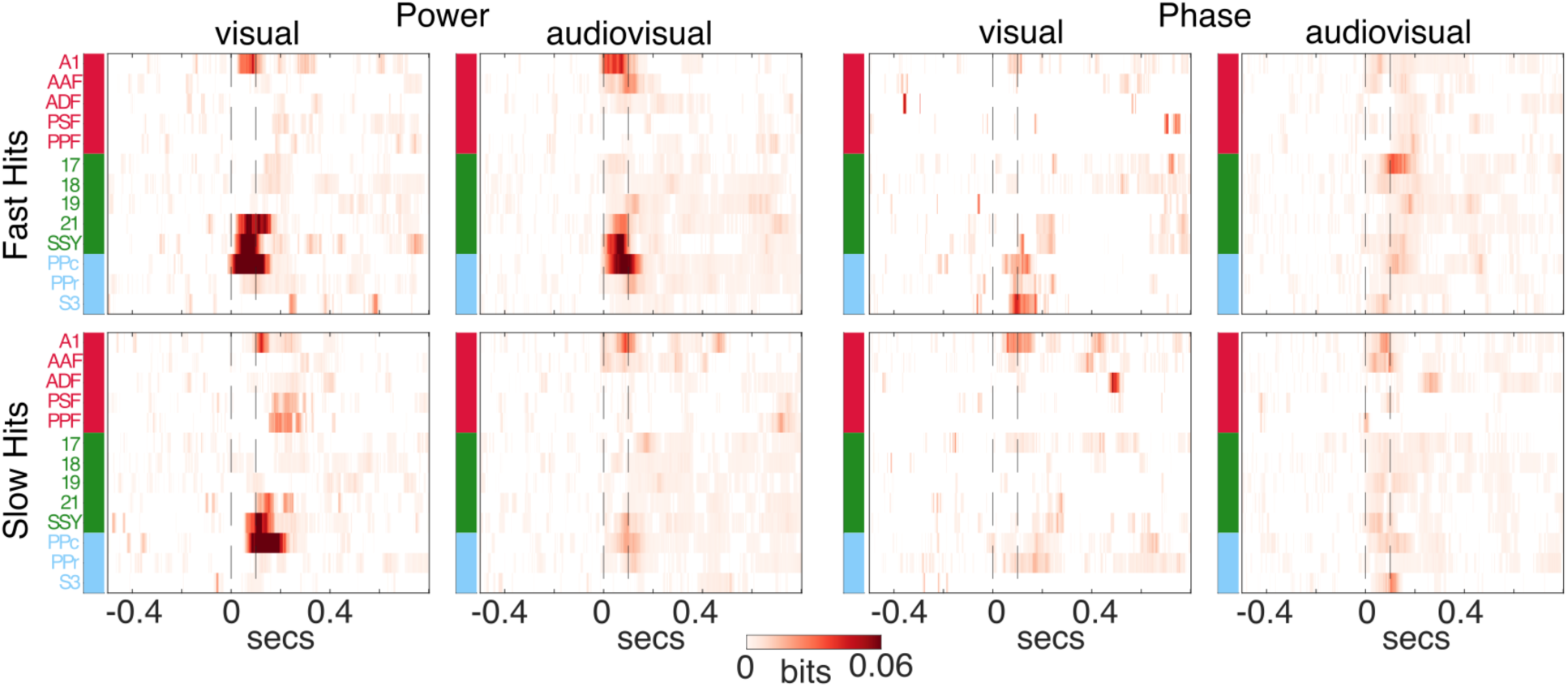
Cortical distribution of alpha-band unique information in fast and slow hit trials for LFP power. Unique information time courses computed separately for fast (top panels) and slow (bottom panels) hit trials. For each brain area, time courses of channels being part of significant pairs of channels (significant joint information) were pooled across animals and averaged. For each channel, we first identified significant time points. Specifically, we z-scored each time course relative to its pre-stimulus baseline, converted z-scores to one-tailed p-values (testing for increases above baseline), and controlled for multiple comparisons across time using the Benjamini–Hochberg FDR procedure (q ≤ 0.05). Only time points surviving FDR correction retained their original values, whereas all non-significant samples were set to zero. We then subtracted from each information time-course its pre-stimulus information averaged across time. Vertical dashed lines mark stimulus onset and offset.

**Figure S5.**
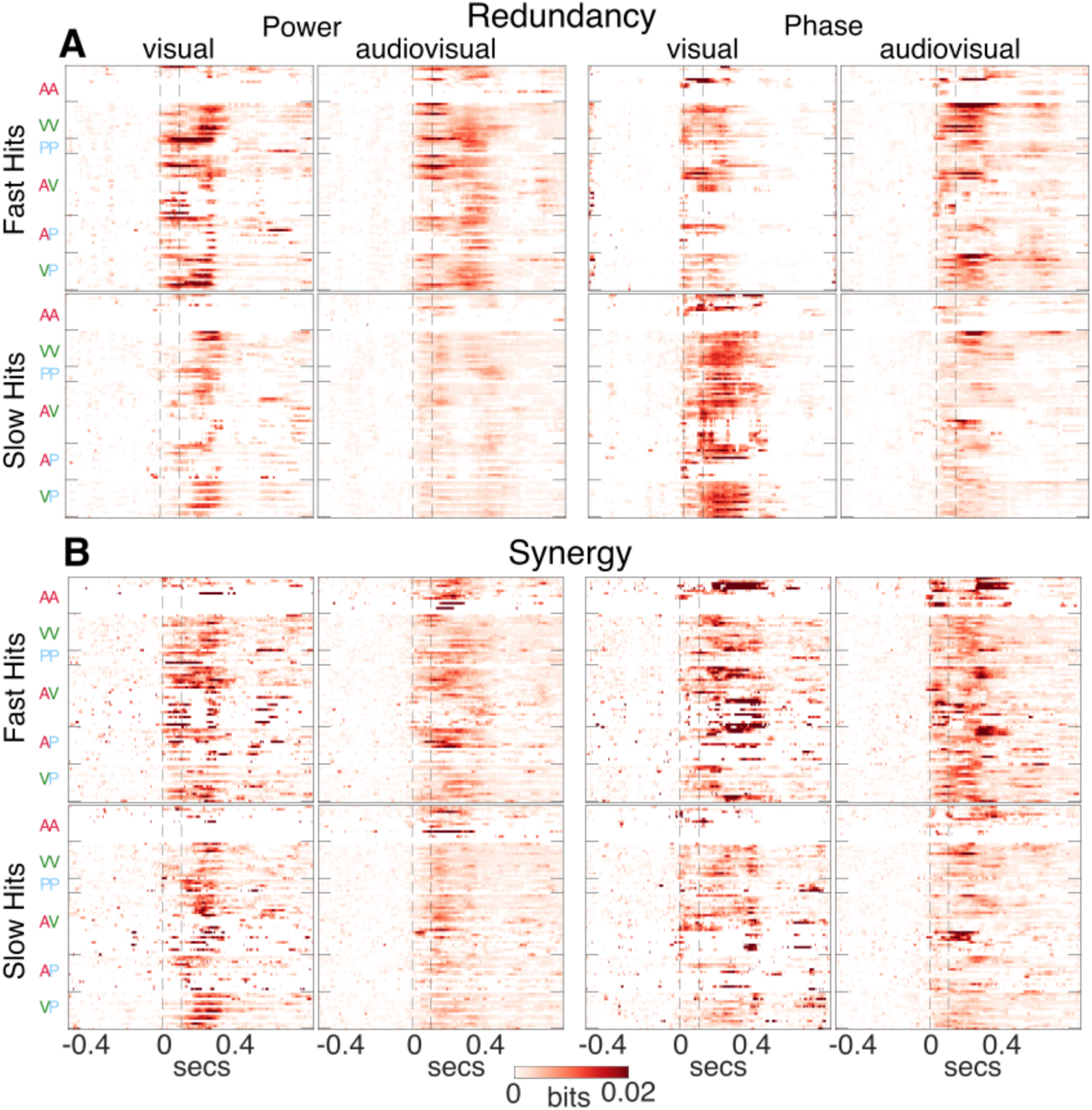
Cortical distribution of theta-band synergistic and redundant information in fast and slow hit trials in LFP power and phase. Heatmaps showing **(A)** redundant and **(B)** synergistic information in significant channel pairs and subjects for each pair of brain areas, grouped by functional system (A, auditory; V, visual; P, parietal). Significant time points in synergistic and redundant components were identified by z-scoring relative to the pre-stimulus baseline and applying Benjamini–Hochberg FDR correction (q ≤ 0.05); non-significant samples were set to zero. Vertical dashed lines denote stimulus onset and offset.

**Figure S6.**
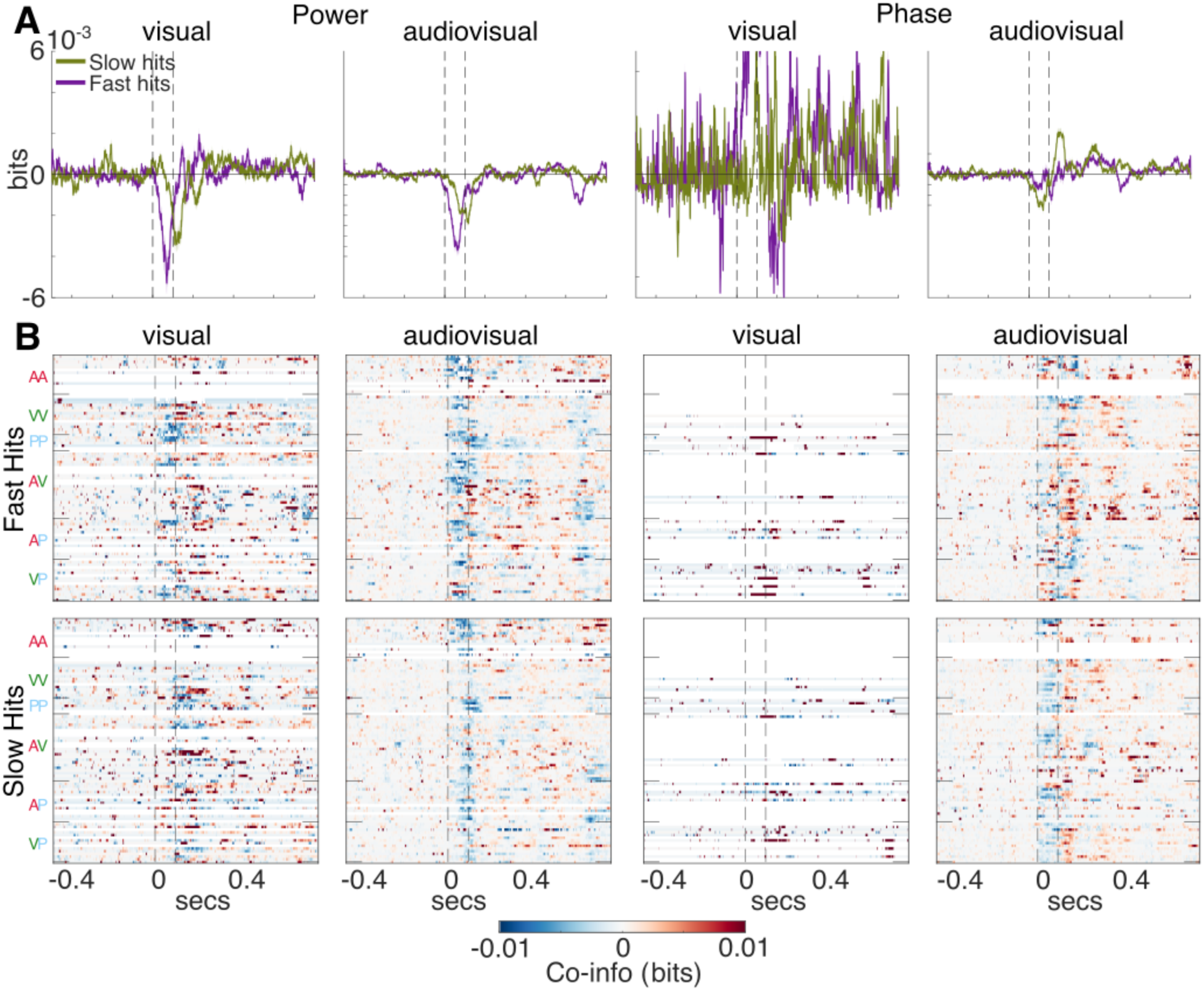
Alpha-band co-information dynamics during fast and slow hit trials. **(A)** Average co-information (synergy minus redundancy) time courses (mean ± SEM) across all channel pairs carrying significant joint information, computed separately for fast and slow hits. **(B)** Heatmaps showing co-information averaged across significant channel pairs and subjects for each pair of brain areas, grouped by functional system (A, auditory; V, visual; P, parietal). Significant time points in synergistic and redundant components were identified by z-scoring relative to the pre-stimulus baseline and applying Benjamini–Hochberg FDR correction (q ≤ 0.05); non-significant samples were set to zero. Vertical dashed lines denote stimulus onset and offset.

**Figure S7.**
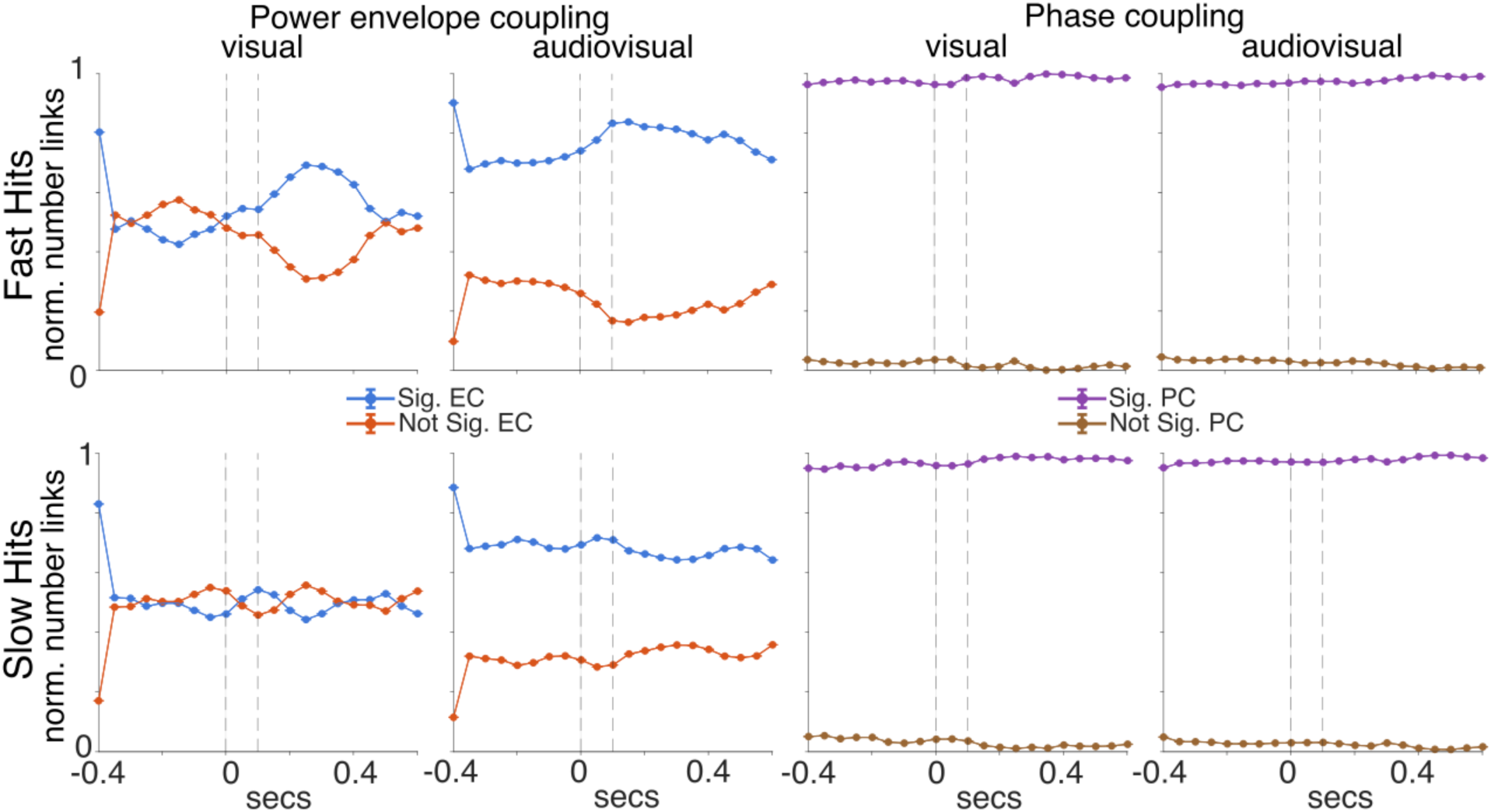
Number of functional links in the two groups of functionally connected and unconnected pairs of channels. Number of functional links (normalized to the total number of links) pooled across subjects categorized as significant and not significant based on a circular-shift permutation test (see Methods). Vertical dashed lines indicate stimulus onset and offset.

**Figure S8.**
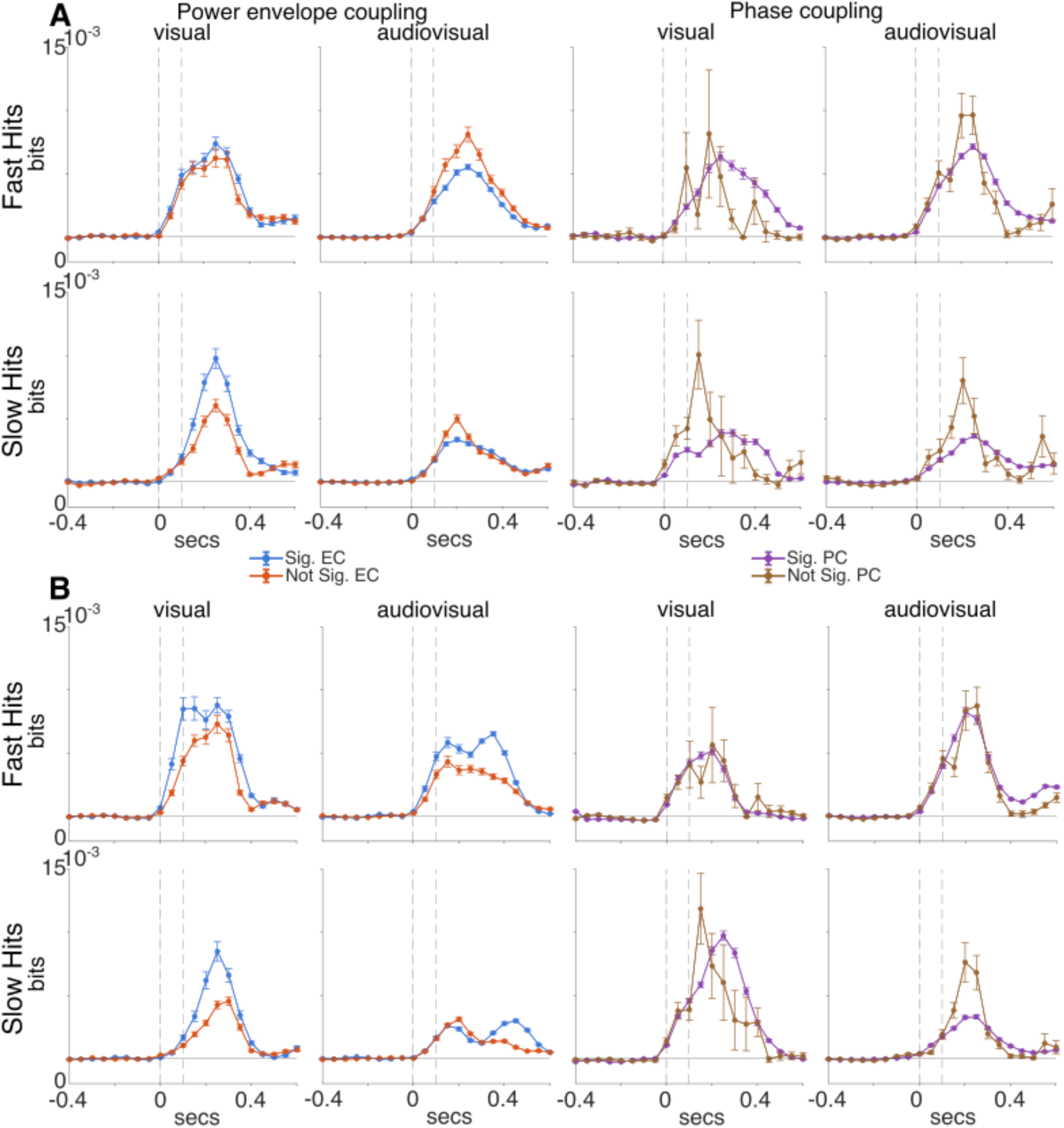
Theta-band synergy and redundancy in functionally coupled channel pairs during fast and slow hits. Time-courses of theta-band synergy (A) and redundancy (B) for channel pairs carrying significant joint information (mean ± SEM), computed separately for functionally connected (Sig EC, Sig PC) and non-connected pairs during fast and slow hits. Co-information was estimated using a sliding window of 100 ms advanced in 50 ms steps. Functional connectivity was assessed within each window using envelope coupling for LFP power and phase coupling for LFP phase, with significance determined by a circular-shift permutation test (p < 0.05). Statistical differences between conditions were evaluated using an unpaired t-test (**p < 0.001). Vertical dashed lines indicate stimulus onset and offset.

**Figure S9.**
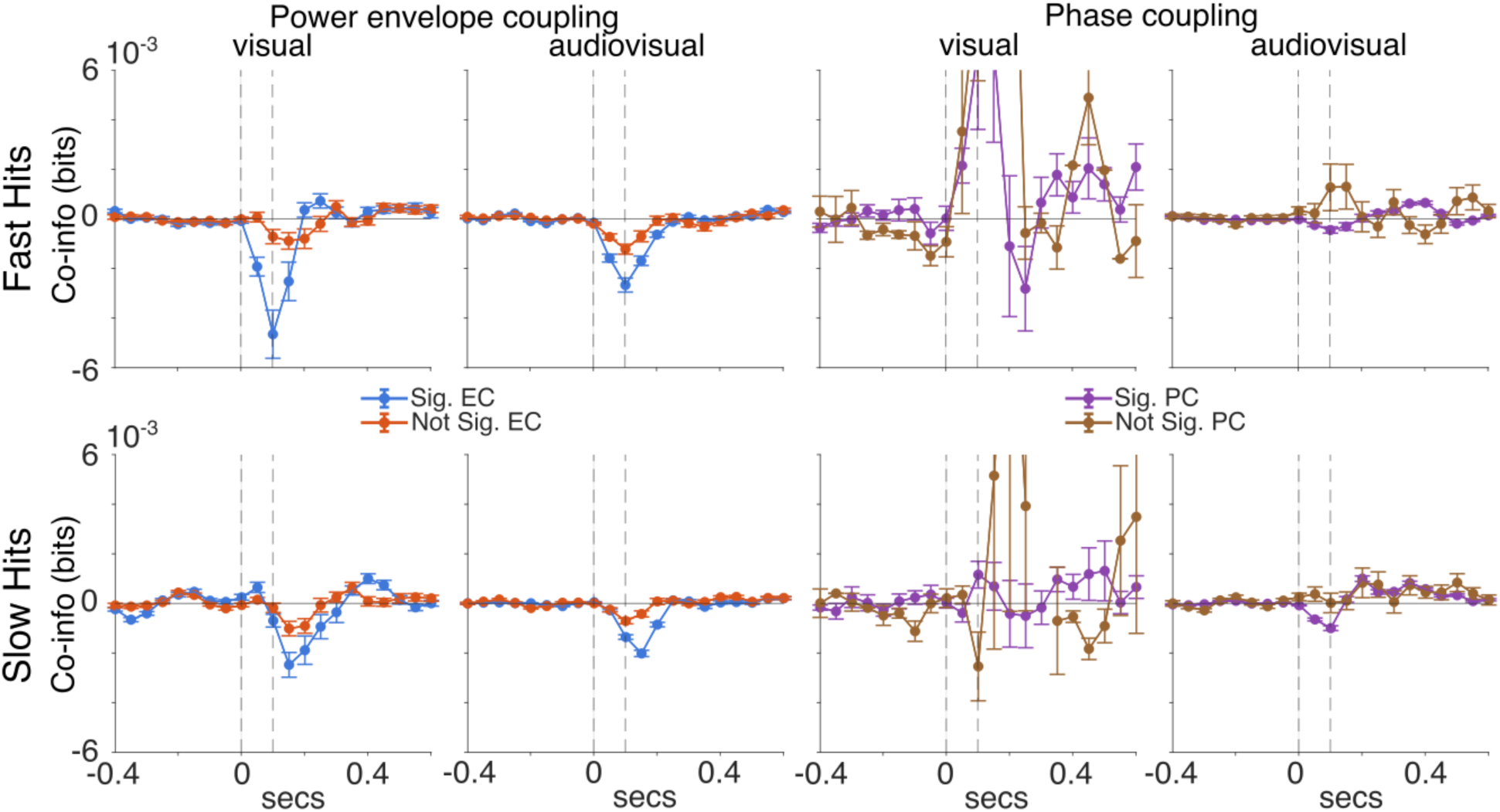
Alpha-band co-information in functionally coupled channel pairs during fast and slow hits. Time courses of alpha-band co-information for channel pairs carrying significant joint information, computed separately for functionally connected (Sig FC) and non-connected pairs during fast and slow hits. Co-information was estimated using a sliding window of 100 ms advanced in 50 ms steps. Functional connectivity was assessed within each window using envelope coupling for LFP power and phase coupling for LFP phase, with significance determined by a circular-shift permutation test (p < 0.05). Statistical differences between conditions were evaluated using an unpaired t-test (**p < 0.001). Vertical dashed lines indicate stimulus onset and offset.

**Figure S10.**
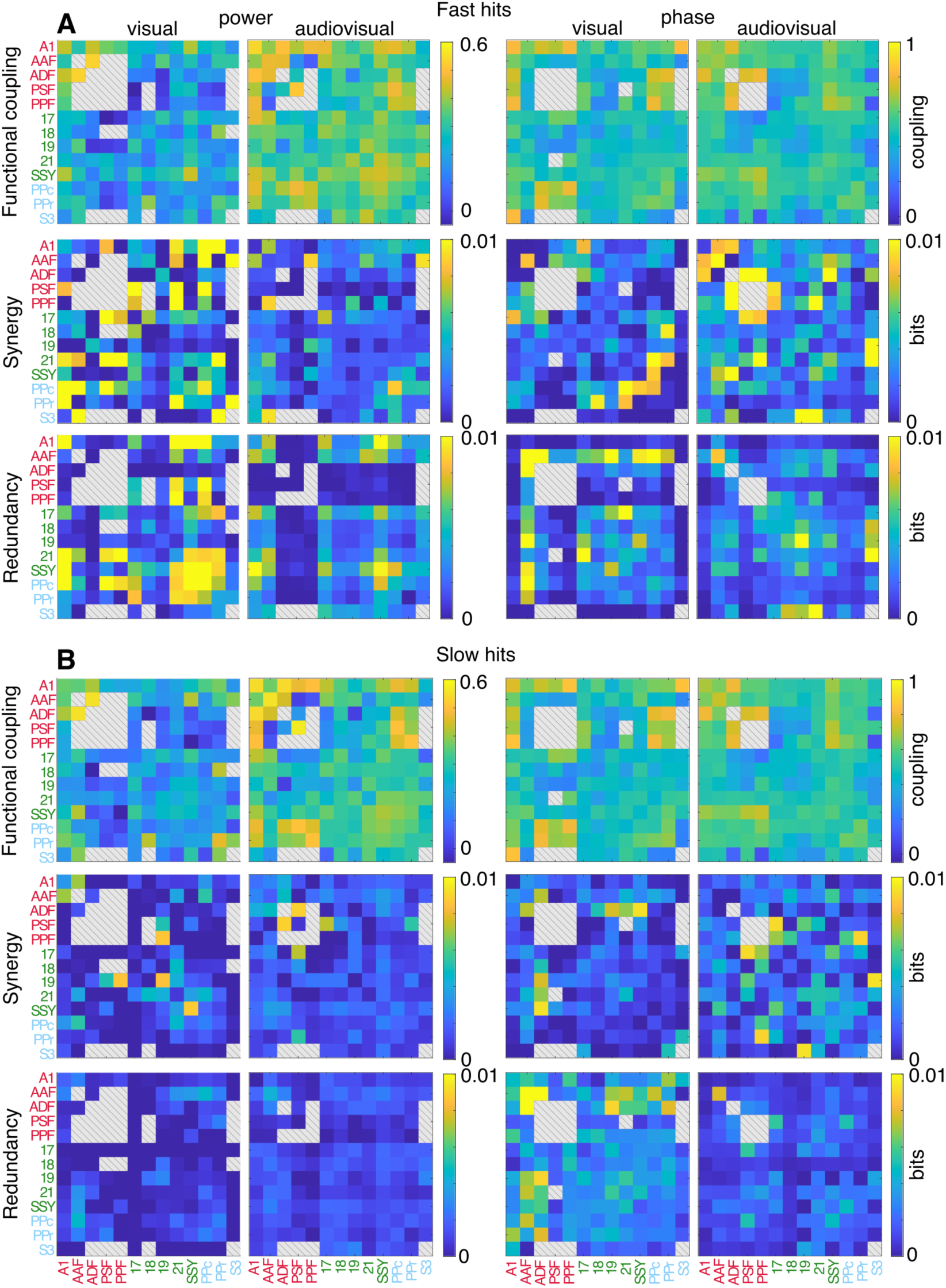
Networks of functional connectivity, synergy and redundancy in the theta band during the stimulus interval. Theta-band synergy and redundancy networks for visual and audiovisual stimuli, averaged within the 100-ms stimulus window for **(A)** fast and **(B)** slow hit trials. The top panels in (A) and (B) show FC matrices for envelope coupling (left) and phase coupling (right). Gray areas with diagonal lines indicate pairs of channels that did not show significant joint information.

**Figure S11.**
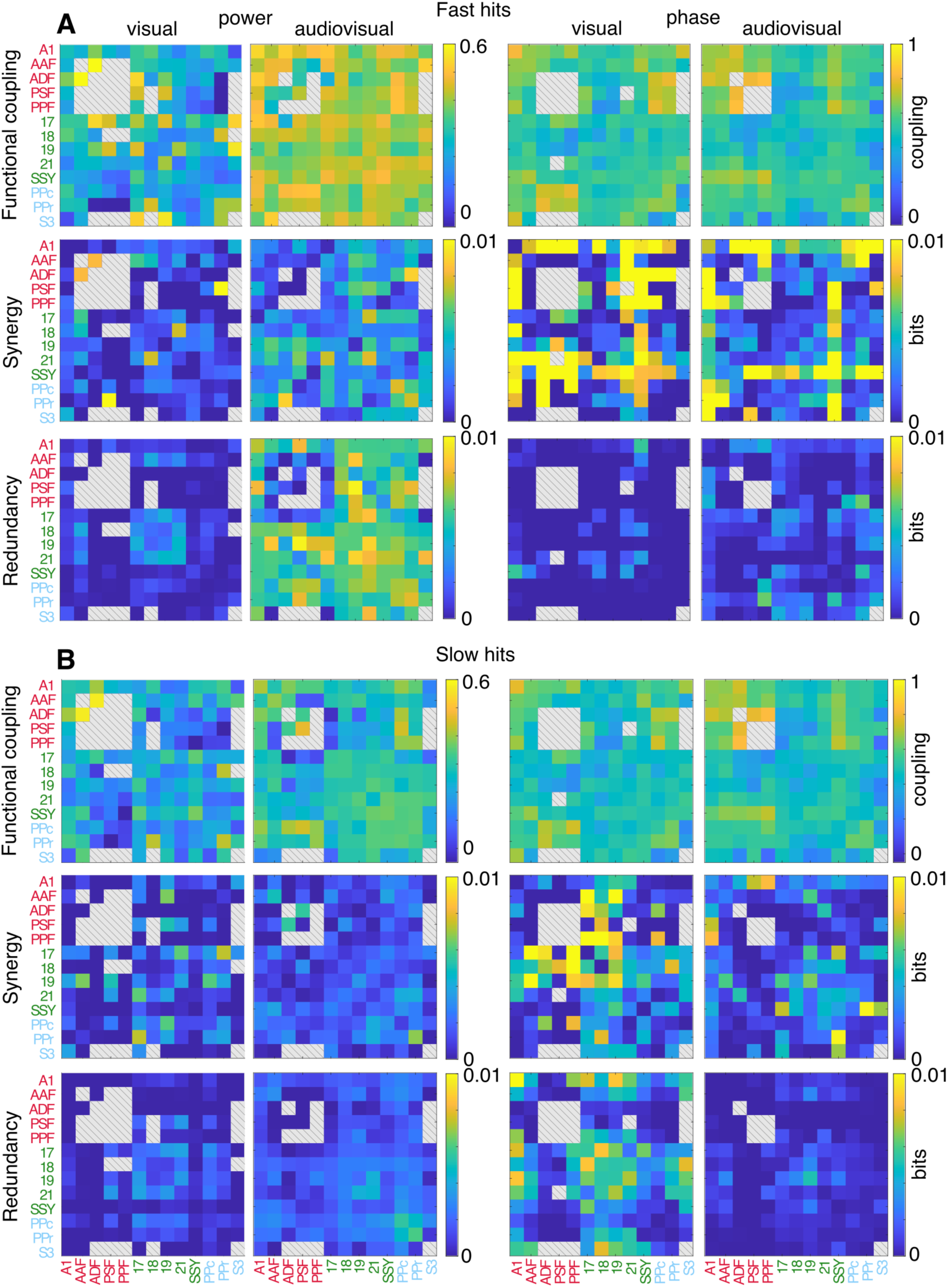
Networks of functional connectivity, synergy and redundancy in the theta band during the post stimulus interval. Theta-band synergy and redundancy networks for visual and audiovisual stimuli, averaged within the 300-400 ms stimulus window for **(A)** fast and **(B)** slow hit trials. The top panels in (A) and (B) show FC matrices for envelope coupling (left) and phase coupling (right). Gray areas with diagonal lines indicate pairs of channels that did not show significant joint information.

**Figure S12.**
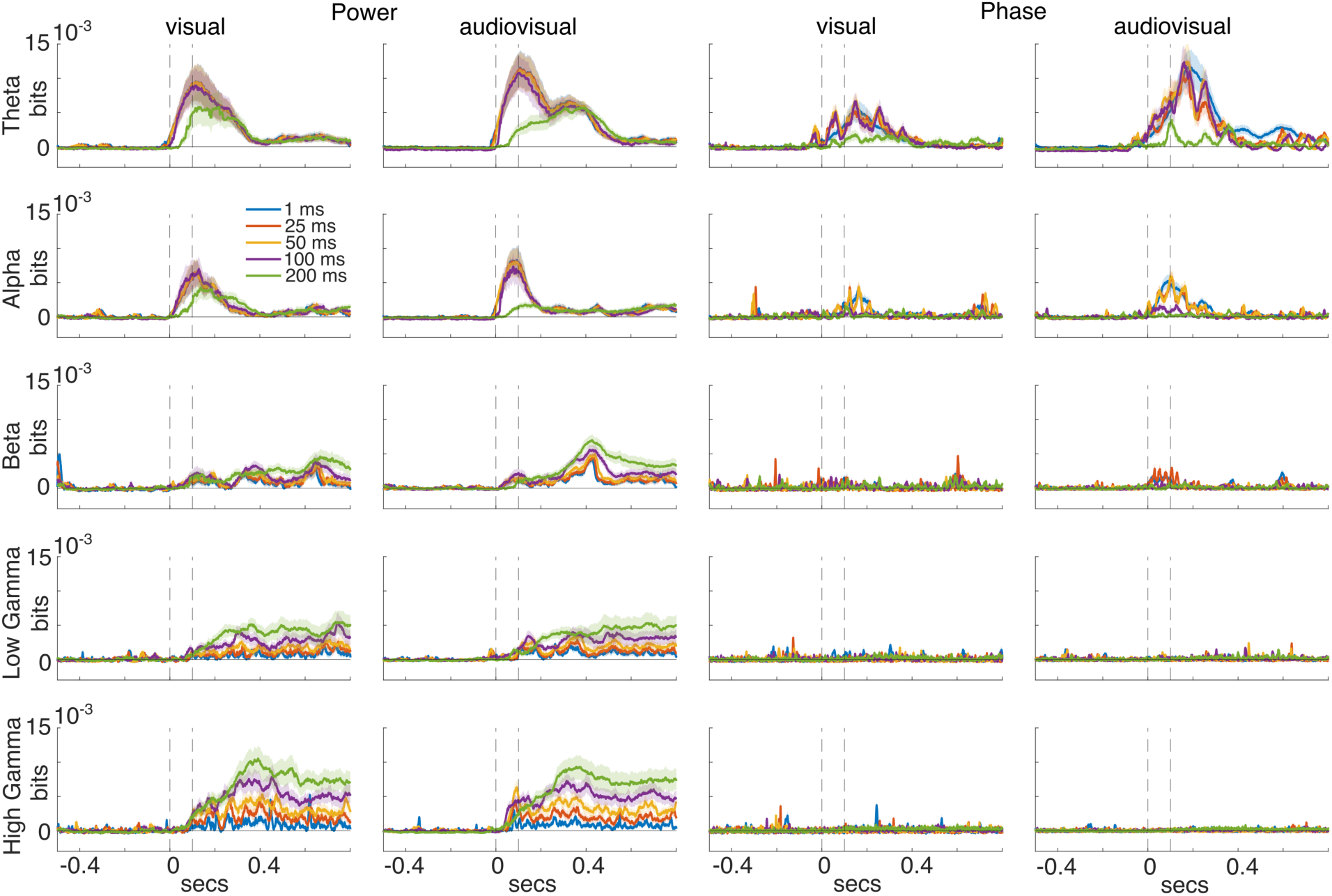
Stimulus-side information computed for different sliding windows. Stimulus information time courses (mean ± SEM) averaged across all 64 ECoG channels for each frequency band. We followed the same approach as described in Fig.2A but changed the window sizes (blue lines = 1ms, green lines = 200 ms) in which we averaged the LFP signals before computing stimulus information. Vertical dashed lines indicate stimulus onset and offset.

**Figure S13.**
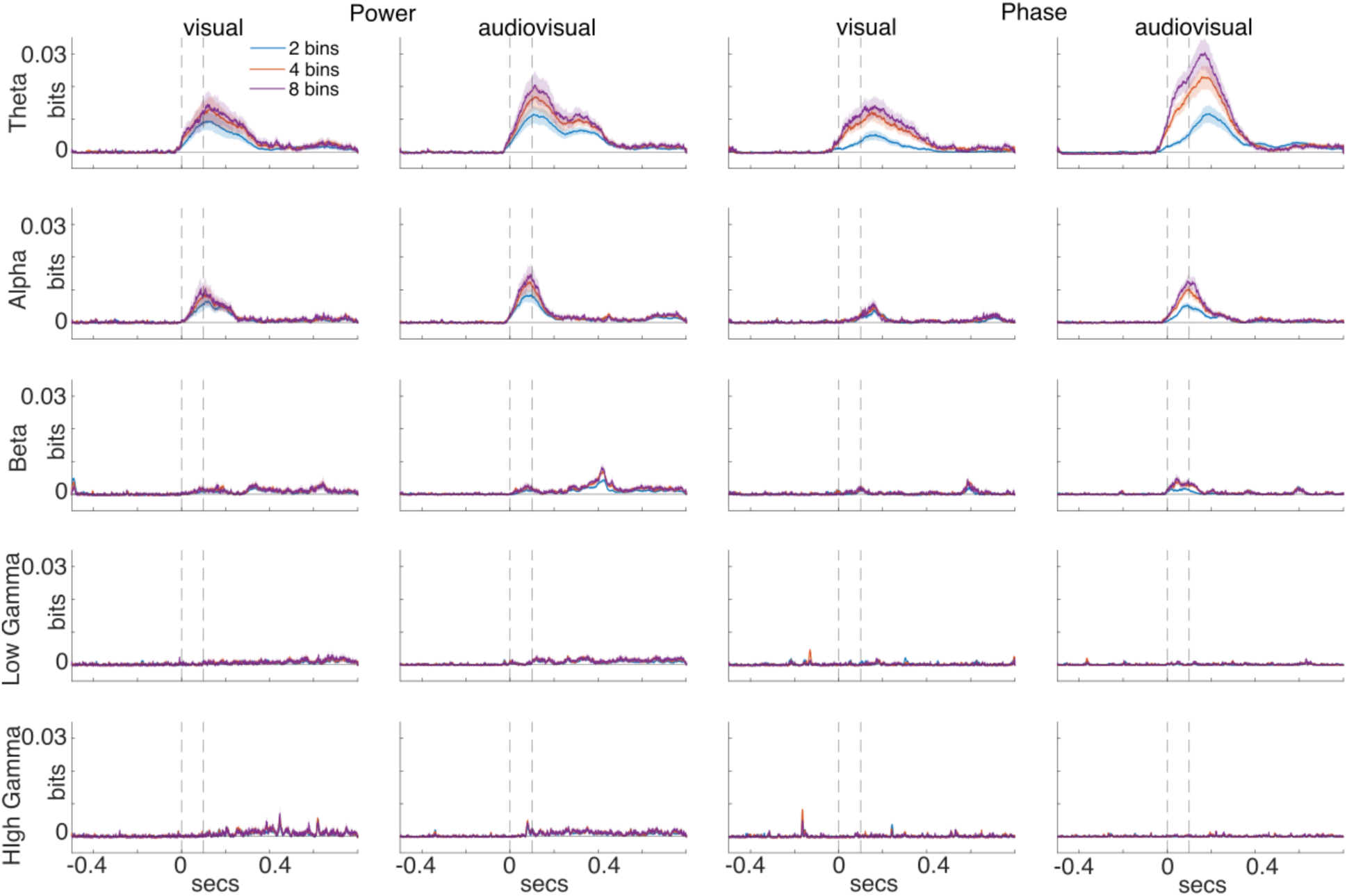
Stimulus-side information for different bin numbers. Stimulus information time courses (mean ± SEM) averaged across all 64 ECoG channels for each frequency band. We followed the same approach as described in Fig.2A but changed the bin number to discretize the LFP signals before computing stimulus information (see Methods). Significant time points were identified by z-scoring single-channel time courses relative to the pre-stimulus baseline and applying Benjamini–Hochberg FDR correction across time (q ≤ 0.05); non-significant samples were set to zero, and pre-stimulus baseline information was subtracted. Vertical dashed lines denote stimulus onset and offset.

## References

1. Attneave, F., 1954. Some informational aspects of visual perception. Psychol. Rev. 61, 183–193. 10.1037/h0054663

2. Averbeck, B.B., Latham, P.E., Pouget, A., 2006. Neural correlations, population coding and computation. Nat. Rev. Neurosci. 7, 358–366. 10.1038/nrn1888

3. Barlow, H.B., 1961. Possible principles underlying the transformation of sensory messages, in: Sensory Communication. MIT Press, pp. 217–233.

4. Bastos, A.M., Vezoli, J., Bosman, C.A., Schoffelen, J.-M., Oostenveld, R., Dowdall, J.R., De Weerd, P., Kennedy, H., Fries, P., 2015a. Visual areas exert feedforward and feedback influences through distinct frequency channels. Neuron 85, 390–401. 10.1016/j.neuron.2014.12.018

5. Bastos, A.M., Vezoli, J., Fries, P., 2015b. Communication through coherence with inter-areal delays. Curr. Opin. Neurobiol. 31, 173–180. 10.1016/j.conb.2014.11.001

6. Belitski, A., Gretton, A., Magri, C., Murayama, Y., Montemurro, M.A., Logothetis, N.K., Panzeri, S., 2008. Low-frequency local field potentials and spikes in primary visual cortex convey independent visual information. J. Neurosci. 28, 5696–5709. 10.1523/JNEUROSCI.0009-08.2008

7. Belitski, A., Panzeri, S., Magri, C., Logothetis, N.K., Kayser, C., 2010. Sensory information in local field potentials and spikes from visual and auditory cortices: time scales and frequency bands. J. Comput. Neurosci. 29, 533–545. 10.1007/s10827-010-0230-y

8. Bertschinger, N., Rauh, J., Olbrich, E., Jost, J., Ay, N., 2014. Quantifying unique information. Entropy 16, 2161–2183. 10.3390/e16042161

9. Bizley, J.K., Nodal, F.R., Bajo, V.M., Nelken, I., King, A.J., 2007. Physiological and anatomical evidence for multisensory interactions in auditory cortex. Cereb. Cortex 17, 2172–2189. 10.1093/cercor/bhl128

10. Boashash, B., 1992. Estimating and interpreting the instantaneous frequency of a signal. I. Fundamentals. Proc. IEEE 80, 520–538. 10.1109/5.135376

11. Brookes, M.J., Woolrich, M., Luckhoo, H., Price, D., Hale, J.R., Stephenson, M.C., Barnes, G.R., Smith, S.M., Morris, P.G., 2011. Investigating the electrophysiological basis of resting state networks using magnetoencephalography. Proc. Natl. Acad. Sci. 108, 16783–16788. 10.1073/pnas.1112685108

12. Cohen, M.R., Maunsell, J.H.R., 2009. Attention improves performance primarily by reducing interneuronal correlations. Nat. Neurosci. 12, 1594–1600. 10.1038/nn.2439

13. Combrisson, E., Basanisi, R., Gueguen, M.C., Rheims, S., Kahane, P., Bastin, J., Brovelli, A., 2024. Neural interactions in the human frontal cortex dissociate reward and punishment learning. eLife 12, RP92938. 10.7554/eLife.92938

14. Combrisson, E., Basanisi, R., Neri, M., Auzias, G., Petri, G., Marinazzo, D., Panzeri, S., Brovelli, A., 2025. Higher-order and distributed synergistic functional interactions encode information gain in goal-directed learning. Nat. Commun. 16, 7179. 10.1038/s41467-025-62507-1

15. Cover, T.M., Thomas, J.A., 2006. Elements of Information Theory, 2nd ed. Wiley-Interscience.

16. Delis, I., Ince, R.A.A., Sajda, P., Wang, Q., 2022. Neural encoding of active multi-sensing enhances perceptual decision-making via a synergistic cross-modal interaction. J. Neurosci. 42, 2344–2355. 10.1523/JNEUROSCI.0861-21.2022

17. Engel, A.K., Fries, P., Singer, W., 2001. Dynamic predictions: oscillations and synchrony in top– down processing. Nat. Rev. Neurosci. 2, 704–716. 10.1038/35094565

18. Engel, A.K., Gerloff, C., 2022. Dynamic functional connectivity: causative or epiphenomenal? Trends Cogn. Sci. 26, 1020–1022. 10.1016/j.tics.2022.09.021

19. Engel, A.K., Gerloff, C., Hilgetag, C.C., Nolte, G., 2013. Intrinsic coupling modes: multiscale interactions in ongoing brain activity. Neuron 80, 867–886. 10.1016/j.neuron.2013.09.038

20. Fiebelkorn, I.C., Kastner, S., 2019. A rhythmic theory of attention. Trends Cogn. Sci. 23, 87–101. 10.1016/j.tics.2018.11.009

21. Francis, N.A., Mukherjee, S., Koçillari, L., Panzeri, S., Babadi, B., Kanold, P.O., 2022. Sequential transmission of task-relevant information in cortical neuronal networks. Cell Rep. 39, 110878. 10.1016/j.celrep.2022.110878

22. Fries, P., 2015. Rhythms for cognition: communication through coherence. Neuron 88, 220–235. 10.1016/j.neuron.2015.09.034

23. Galindo-Leon, E.E., Hollensteiner, K.J., Pieper, F., Engler, G., Nolte, G., Engel, A.K., 2025a. Dynamic changes in large-scale functional connectivity prior to stimulation determine performance in a multisensory task. Front. Syst. Neurosci. 19, 1524547. 10.3389/fnsys.2025.1524547

24. Galindo-Leon, E.E., Nolte, G., Pieper, F., Engler, G., Engel, A.K., 2025b. Causal interactions between amplitude correlation and phase coupling in cortical networks. Sci. Rep. 15, 11975. 10.1038/s41598-025-95306-1

25. Galindo-Leon, E.E., Stitt, I., Pieper, F., Stieglitz, T., Engler, G., Engel, A.K., 2019. Context-specific modulation of intrinsic coupling modes shapes multisensory processing. Sci. Adv. 5, eaar7633. 10.1126/sciadv.aar7633

26. Gelens, F., Äijälä, J., Roberts, L., Komatsu, M., Uran, C., Jensen, M.A., Miller, K.J., Ince, R.A.A., Garagnani, M., Vinck, M., Canales-Johnson, A., 2024. Distributed representations of prediction error signals across the cortical hierarchy are synergistic. Nat. Commun. 15, 3941. 10.1038/s41467-024-48329-7

27. Giraud, A.-L., Poeppel, D., 2012. Cortical oscillations and speech processing: emerging computational principles and operations. Nat. Neurosci. 15, 511–517. 10.1038/nn.3063

28. Gray, C.M., König, P., Engel, A.K., Singer, W., 1989. Oscillatory responses in cat visual cortex exhibit inter-columnar synchronization which reflects global stimulus properties. Nature 338, 334–337. 10.1038/338334a0

29. Greco, A., Moser, J., Preissl, H., Siegel, M., 2024. Predictive learning shapes the representational geometry of the human brain. Nat. Commun. 15, 9670. 10.1038/s41467-024-54032-4

30. Gross, J., Hoogenboom, N., Thut, G., Schyns, P., Panzeri, S., Belin, P., Garrod, S., 2013. Speech rhythms and multiplexed oscillatory sensory coding in the human brain. PLOS Biol. 11, e1001752. 10.1371/journal.pbio.1001752

31. Helfrich, R.F., Fiebelkorn, I.C., Szczepanski, S.M., Lin, J.J., Parvizi, J., Knight, R.T., Kastner, S., 2018. Neural mechanisms of sustained attention are rhythmic. Neuron 99, 854–865.e5. 10.1016/j.neuron.2018.07.032

32. Hernández, A., Nácher, V., Luna, R., Zainos, A., Lemus, L., Alvarez, M., Vázquez, Y., Camarillo, L., Romo, R., 2010. Decoding a perceptual decision process across cortex. Neuron 66, 300–314. 10.1016/j.neuron.2010.03.031

33. Hipp, J.F., Hawellek, D.J., Corbetta, M., Siegel, M., Engel, A.K., 2012. Large-scale cortical correlation structure of spontaneous oscillatory activity. Nat. Neurosci. 15, 884–890. 10.1038/nn.3101

34. Hollensteiner, K.J., Pieper, F., Engler, G., König, P., Engel, A.K., 2015. Crossmodal integration improves sensory detection thresholds in the ferret. PLOS ONE 10, e0124952. 10.1371/journal.pone.0124952

35. International Brain Laboratory, et al., 2025. A brain-wide map of neural activity during complex behaviour. Nature 645, 177–191. 10.1038/s41586-025-09235-0

36. Kayser, C., Montemurro, M.A., Logothetis, N.K., Panzeri, S., 2009. Spike-phase coding boosts and stabilizes information carried by spatial and temporal spike patterns. Neuron 61, 597– 608. 10.1016/j.neuron.2009.01.008

37. Khilkevich, A., Lohse, M., Low, R., Orsolic, I., Bozic, T., Windmill, P., Mrsic-Flogel, T.D., 2024. Brain-wide dynamics linking sensation to action during decision-making. Nature 634, 890– 900. 10.1038/s41586-024-07908-w

38. Koçillari, L., Celotto, M., Francis, N.A., Mukherjee, S., Babadi, B., Kanold, P.O., Panzeri, S., 2023. Behavioural relevance of redundant and synergistic stimulus information between functionally connected neurons in mouse auditory cortex. Brain Inform. 10, 34. 10.1186/s40708-023-00212-9

39. Lachaux, J.-P., Rodriguez, E., Martinerie, J., Varela, F.J., 1999. Measuring phase synchrony in brain signals. Hum. Brain Mapp. 8, 194–208. 10.1002/(sici)1097-0193(1999)8:4%3C194::aid-hbm4%3E3.0.co;2-c

40. Lam, N.H., Mukherjee, A., Wimmer, R.D., Nassar, M.R., Chen, Z.S., Halassa, M.M., 2025. Prefrontal transthalamic uncertainty processing drives flexible switching. Nature 637, 127–136. 10.1038/s41586-024-08180-8

41. Laughlin, S., 1981. A simple coding procedure enhances a neuron’s information capacity. Z. Naturforschung C Biosci. 36, 910–912. 10.1515/znc-1981-9-1040

42. Lemke, S.M., Celotto, M., Maffulli, R., Ganguly, K., Panzeri, S., 2024. Information flow between motor cortex and striatum reverses during skill learning. Curr. Biol. 34, 1831–1843.e7. 10.1016/j.cub.2024.03.023

43. Liebe, S., Hoerzer, G.M., Logothetis, N.K., Rainer, G., 2012. Theta coupling between V4 and prefrontal cortex predicts visual short-term memory performance. Nat. Neurosci. 15, 456–462. 10.1038/nn.3038

44. Lorenz, G.M., Engel, N.M., Celotto, M., Koçillari, L., Curreli, S., Fellin, T., Panzeri, S., 2025. MINT: A toolbox for the analysis of multivariate neural information coding and transmission. PLOS Comput. Biol. 21, e1012934. 10.1371/journal.pcbi.1012934

45. Lorenz, G.M., Engel, N.M., Koçillari, L., Celotto, M., Orsenigo, D., Curreli, S., Malerba, S.B., Engel, A.K., Kayser, C., Fellin, T., Luppi, A.I., Panzeri, S., 2026. Sampling bias corrections for discrete and Gaussian partial information decompositions. Patterns. 10.1016/j.patter.2026.101619

46. Louviot, S., Radanovic, A., Martin, I., Patchell, A., Alkhoury, L., Scanavini, G., Schiff, N.D., Hill, N.J., Shah, S.A., 2025. Cortical oscillatory dynamics underlying response speed: insights from high-density EEG and the attention network test. Cereb. Cortex 35, bhaf316. 10.1093/cercor/bhaf316

47. Luppi, A.I., Mediano, P.A.M., Rosas, F.E., Allanson, J., Pickard, J., Carhart-Harris, R.L., Williams, G.B., Craig, M.M., Finoia, P., Owen, A.M., Naci, L., Menon, D.K., Bor, D., Stamatakis, E.A., 2024a. A synergistic workspace for human consciousness revealed by Integrated Information Decomposition. eLife 12, RP88173. 10.7554/eLife.88173

48. Luppi, A.I., Mediano, P.A.M., Rosas, F.E., Holland, N., Fryer, T.D., O’Brien, J.T., Rowe, J.B., Menon, D.K., Bor, D., Stamatakis, E.A., 2022. A synergistic core for human brain evolution and cognition. Nat. Neurosci. 25, 771–782. 10.1038/s41593-022-01070-0

49. Luppi, A.I., Rosas, F.E., Mediano, P.A.M., Menon, D.K., Stamatakis, E.A., 2024b. Information decomposition and the informational architecture of the brain. Trends Cogn. Sci. 28, 352–368. 10.1016/j.tics.2023.11.005

50. Michalareas, G., Vezoli, J., van Pelt, S., Schoffelen, J.-M., Kennedy, H., Fries, P., 2016. Alpha-beta and gamma rhythms subserve feedback and feedforward influences among human visual cortical areas. Neuron 89, 384–397. 10.1016/j.neuron.2015.12.018

51. Montemurro, M.A., Rasch, M.J., Murayama, Y., Logothetis, N.K., Panzeri, S., 2008. Phase-of-firing coding of natural visual stimuli in primary visual cortex. Curr. Biol. 18, 375–380. 10.1016/j.cub.2008.02.023

52. Nolte, G., Bai, O., Wheaton, L., Mari, Z., Vorbach, S., Hallett, M., 2004. Identifying true brain interaction from EEG data using the imaginary part of coherency. Clin. Neurophysiol. 115, 2292–2307. 10.1016/j.clinph.2004.04.029

53. Nolte, G., Galindo-Leon, E.E., Li, Z., Liu, X., Engel, A.K., 2020. Mathematical relations between measures of brain connectivity estimated from electrophysiological recordings for gaussian distributed data. Front. Neurosci. 14. 10.3389/fnins.2020.577574

54. Panzeri, S., Brunel, N., Logothetis, N.K., Kayser, C., 2010. Sensory neural codes using multiplexed temporal scales. Trends Neurosci. 33, 111–120. 10.1016/j.tins.2009.12.001

55. Panzeri, S., Moroni, M., Safaai, H., Harvey, C.D., 2022. The structures and functions of correlations in neural population codes. Nat. Rev. Neurosci. 23, 551–567. 10.1038/s41583-022-00606-4

56. Park, H., Ince, R.A.A., Schyns, P.G., Thut, G., Gross, J., 2018. Representational interactions during audiovisual speech entrainment: Redundancy in left posterior superior temporal gyrus and synergy in left motor cortex. PLOS Biol. 16, e2006558. 10.1371/journal.pbio.2006558

57. Pesaran, B., Nelson, M.J., Andersen, R.A., 2008. Free choice activates a decision circuit between frontal and parietal cortex. Nature 453, 406–409. 10.1038/nature06849

58. Pope, M., Varley, T.F., Puxeddu, M.G., Faskowitz, J., Sporns, O., 2025. Time-varying synergy/redundancy dominance in the human cerebral cortex. J. Phys. Complex. 6, 015015. 10.1088/2632-072X/adbaa9

59. Proca, A.M., Rosas, F.E., Luppi, A.I., Bor, D., Crosby, M., Mediano, P.A.M., 2024. Synergistic information supports modality integration and flexible learning in neural networks solving multiple tasks. PLOS Comput. Biol. 20, e1012178. 10.1371/journal.pcbi.1012178

60. Quian Quiroga, R., Panzeri, S., 2009. Extracting information from neuronal populations: information theory and decoding approaches. Nat. Rev. Neurosci. 10, 173–185.

61. Reid, A.T., Headley, D.B., Mill, R.D., Sanchez-Romero, R., Uddin, L.Q., Marinazzo, D., Lurie, D.J., Valdés-Sosa, P.A., Hanson, S.J., Biswal, B.B., Calhoun, V., Poldrack, R.A., Cole, M.W., 2019. Advancing functional connectivity research from association to causation. Nat. Neurosci. 22, 1751–1760. 10.1038/s41593-019-0510-4

62. Roberts, L., Äijälä, J., Burger, F., Uran, C., Jensen, M.A., Miller, K.J., Ince, R.A.A., Vinck, M., Hermes, D., Canales-Johnson, A., 2026. Broadband synergy versus oscillatory redundancy in the visual cortex. Nat. Commun. 17, 5568. 10.1038/s41467-026-72444-2

63. Schroeder, C.E., Lakatos, P., 2009. Low-frequency neuronal oscillations as instruments of sensory selection. Trends Neurosci. 32, 9–18. 10.1016/j.tins.2008.09.012

64. Shannon, C.E., 1948. A Mathematical Theory of Communication. Bell Syst. Tech. J. 27, 379–423. 10.1002/j.1538-7305.1948.tb01338.x

65. Siegel, M., Buschman, T.J., Miller, E.K., 2015. Cortical information flow during flexible sensorimotor decisions. Science 348, 1352–1355. 10.1126/science.aab0551

66. Siegel, M., Donner, T.H., Engel, A.K., 2012. Spectral fingerprints of large-scale neuronal interactions. Nat. Rev. Neurosci. 13, 121–134. 10.1038/nrn3137

67. Siems, M., Siegel, M., 2020. Dissociated neuronal phase– and amplitude-coupling patterns in the human brain. NeuroImage 209, 116538. 10.1016/j.neuroimage.2020.116538

68. Simoncelli, E.P., Olshausen, B.A., 2001. Natural image statistics and neural representation. Annu. Rev. Neurosci. 24, 1193–1216. 10.1146/annurev.neuro.24.1.1193

69. Steinmetz, N.A., Zatka-Haas, P., Carandini, M., Harris, K.D., 2019. Distributed coding of choice, action and engagement across the mouse brain. Nature 576, 266–273. 10.1038/s41586-019-1787-x

70. Valente, M., Pica, G., Bondanelli, G., Moroni, M., Runyan, C.A., Morcos, A.S., Harvey, C.D., Panzeri, S., 2021. Correlations enhance the behavioral readout of neural population activity in association cortex. Nat. Neurosci. 24, 975–986. 10.1038/s41593-021-00845-1

71. Varley, T.F., Pope, M., Faskowitz, J., Sporns, O., 2023a. Multivariate information theory uncovers synergistic subsystems of the human cerebral cortex. Commun. Biol. 6, 451. 10.1038/s42003-023-04843-w

72. Varley, T.F., Pope, M., Puxeddu, M.G., Faskowitz, J., Sporns, O., 2023b. Partial entropy decomposition reveals higher-order information structures in human brain activity. Proc. Natl. Acad. Sci. 120, e2300888120. 10.1073/pnas.2300888120

73. Varley, T.F., Sporns, O., Schaffelhofer, S., Scherberger, H., Dann, B., 2023c. Information-processing dynamics in neural networks of macaque cerebral cortex reflect cognitive state and behavior. Proc. Natl. Acad. Sci. 120, e2207677120. 10.1073/pnas.2207677120

74. Williams, P.L., Beer, R.D., 2010. Nonnegative decomposition of multivariate information. 10.48550/arXiv.1004.2515

75. Wilming, N., Murphy, P.R., Meyniel, F., Donner, T.H., 2020. Large-scale dynamics of perceptual decision information across human cortex. Nat. Commun. 11, 5109. 10.1038/s41467-020-18826-6

76. Womelsdorf, T., Fries, P., Mitra, P.P., Desimone, R., 2006. Gamma-band synchronization in visual cortex predicts speed of change detection. Nature 439, 733–736. 10.1038/nature04258

77. Ye, T., Romero-Sosa, J.L., Rickard, A., Aguirre, C.G., Wikenheiser, A.M., Blair, H.T., Izquierdo, A., 2023. Theta oscillations in anterior cingulate cortex and orbitofrontal cortex differentially modulate accuracy and speed in flexible reward learning. Oxf. Open Neurosci. 2, kvad005. 10.1093/oons/kvad005

